# Novel signaling hub of insulin receptor, dystrophin glycoprotein complex and plakoglobin regulates muscle size

**DOI:** 10.1101/517789

**Authors:** Yara Eid Mutlak, Dina Aweida, Alexandra Volodin, Bar Ayalon, Nitsan Dahan, Anna Parnis, Shenhav Cohen

**Author notes:** Equal contribution (Yara Eid Mutlak, Dina Aweida). Correspondence to Dr. Shenhav Cohen: Faculty of Biology, Technion Institute of Technology, Haifa 32000, Israel.

## Abstract

Signaling through the insulin receptor governs central physiological functions related to cell growth and metabolism. Here we show by tandem native protein complex purification approach and super-resolution STED microscopy that insulin receptor activity requires association with the fundamental structural module in muscle, the dystrophin glycoprotein complex (DGC), and the desmosomal component plakoglobin (γ-catenin). The integrity of this high-molecular-mass assembly renders skeletal muscle susceptibility to insulin because DGC-insulin receptor dissociation by plakoglobin downregulation reduced insulin signaling and caused atrophy. Furthermore, impaired insulin receptor function in muscles from diabetic mice reduced plakoglobin-DGC-insulin receptor content on the plasma membrane; however, plakoglobin overexpression alone restored DGC association with the insulin receptor, and stimulated glucose uptake. Our findings establish DGC as a signaling hub, containing plakoglobin as an auxiliary subunit, and provide a possible mechanism for the insulin resistance in Duchenne Muscular Dystrophy, and for the cardiomyopathies seen with plakoglobin mutations.

---

The reciprocal dynamics between tissue architecture and function is maintained by the integrity of structural complexes and the transmission of growth signals from cell surface receptors. In skeletal muscle, impaired structural integrity or loss of growth-promoting signals lead to reduced cell size and contractile capacity, and different myopathies in human^1,2^. PI3K-Akt signaling is transmitted via the insulin receptor, and reduced activity of this pathway causes muscle atrophy, and largely contributes to the development of insulin resistance in type-2 diabetes^3–6^. A similar reduction in insulin sensitivity^7,8^ is often caused by the dissociation of dystrophin glycoprotein complex (DGC) in Duchenne muscular dystrophy (DMD)^9,10^, although the precise mechanism is unknown. Specifically, it is unclear how the integrity of the pivotal structural unit of skeletal muscle, the DGC, affects insulin-PI3K-Akt signaling. Here, we demonstrate for the first time that the DGC is physically and functionally linked to the insulin receptor on the plasma membrane.

The DGC plays a structural role by anchoring the cytoskeleton to the extracellular matrix (ECM) to provide mechanical support and protect the sarcolemma against stress imposed during muscle contraction^11^. It is composed of dystrophin, dystroglycans, sarcoglycans, sarcospan, dystrobrevins and syntrophin^11,12^, and loss of dystrophin causes Duchenne muscular dystrophy (DMD). In skeletal muscle, the DGC is confined to subsarcolemmal structures called “costameres”, which reside at the sarcolemma in register with the subjacent sarcomeric Z-bands, just where desmin filaments abut the plasma membrane^13^. In addition, DGC seems to regulate signaling pathways that control cell growth and differentiation^12,14,15^. For example, β-dystroglycan association with the ECM protein laminin stimulates PI3K-Akt and MAPK signaling in skeletal muscle^12,16^ and IGF-1 activity in differentiating oligodendrocyte^15^. Conversely, DGC dissociation in muscles from DMD patients often correlates with the development of insulin resistance^7,8^. Furthermore, reduced DGC integrity in tumor-bearing mice induces the expression of atrophy-related genes and promotes muscle wasting^14^. The present studies uncover a novel signaling hub of DGC and insulin receptor, which resides in costameres and whose integrity and function require the desmosomal component plakoglobin.

Plakoglobin is a component of desmosome adhesion complexes that are prominent in tissues that must withstand mechanical stress, especially cardiomyocytes and epithelia^17–20^. In epithelia, plakoglobin regulates signaling pathways (e.g., by Wnt) that control cell motility^21^, growth, and differentiation^22^. We have recently shown that plakoglobin is of prime importance in regulating skeletal muscle size because it binds to the insulin receptor and enhances the activity of the PI3K-Akt pathway^23^. Activation of this pathway by IGF-I or insulin promotes glucose uptake, and overall protein synthesis, and inhibits protein degradation^2,24,25^. In untreated diabetes, sepsis, and cancer cachexia^26^, impaired signaling through this pathway causes severe muscle wasting, which can be inhibited by the activation of PI3K-Akt-FoxO signaling^27^. Because changes in plakoglobin levels alone influence PI3K-Akt-FoxO pathway^23^, perturbation of its function may contribute to insulin resistance in disease. We show here that plakoglobin is of prime importance for maintenance of muscle size by stabilizing the DGC and mediating its association with and activation of the insulin receptor.

## RESULTS

### Isolation of plakoglobin containing-protein complexes from skeletal muscle

To understand the role of plakoglobin in maintenance of normal muscle size, we isolated the complexes that it forms in skeletal muscle. For this purpose, we modified a recently developed approach^28^, combining tandem native purification of protein complexes and mass spectrometry. Insoluble fractions of 1g lower limb mouse muscles (the 10,000 × g pellet) were washed with high ATP buffer to dissociate the myofibrils, and high pH buffer (pH 9 at 37°C) to solubilize cytoskeletal and membrane bound protein complexes (Fig. 1a,b). Following protein concentration with ammonium sulfate precipitation, the re-solubilized pellets were analyzed by gel filtration column (Superdex 200 10/300) (Fig. 1a), and showed two distinct protein peaks of high and low molecular masses (Fig. 1c) containing high and low molecular weight (MW) proteins (Fig. 1d). This elution profile seems to characterize skeletal muscle generally because a similar two-peak pattern has been obtained from the analysis of rat skeletal muscles (**Supplementary Fig. 1**). Plakoglobin was detected in considerable amounts in the high MW protein peak, and thus seems to be a component of multiprotein assemblies (Fig. 1d).

**Figure 1.**
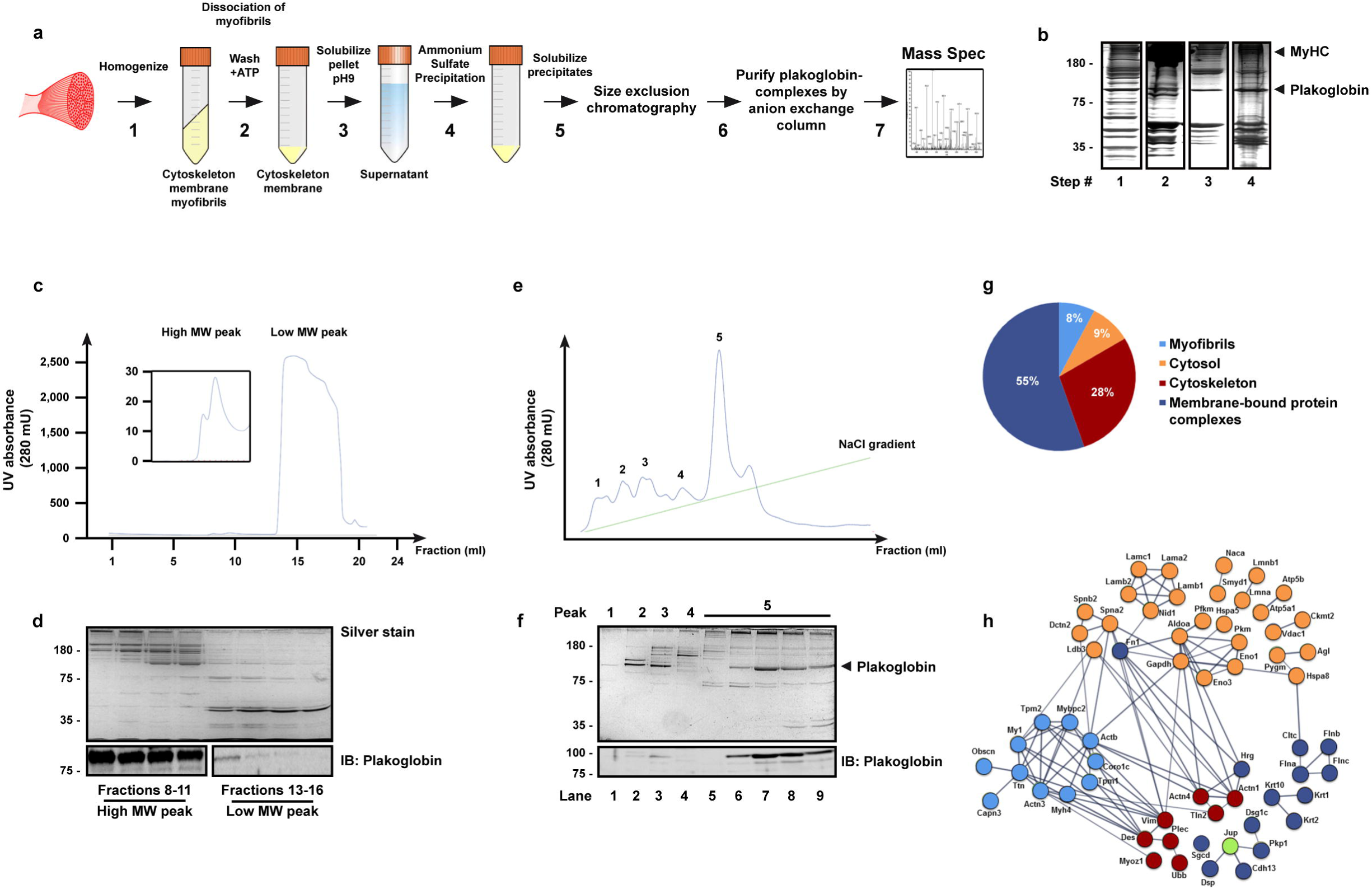
Isolation of plakoglobin-containing protein complexes from mouse skeletal muscle. a. Scheme of the native protein complex purification and proteomic approach.
b. Input from steps 1-4 in **(a)** was analyzed by SDS-PAGE and silver staining (a representative of three independent analyses). Black lines indicate the removal of intervening lanes for clarity and presentation purposes.
c. Size-exclusion chromatography of solubilized ammonium sulfate precipitates from slep 5 in **(a)** yielded two distinct high and low MW protein peaks.
d. SDS-PAGE of fractions from the high (fractions #8-11) and low (fractions #13-16) MW protein peaks in **(c)** followed by silver staining (top panel) or immunoblotting with plakoglobin antibody (lower panel)(a representative of three independent analyses). Black lines indicate the removal of intervening lanes for clarity and presentation purposes.
e. Isolation of plakoglobin containing protein complexes by anion exchange chromatography. Fractions #8-11 from high MW protein peak in **(d)** were pooled and applied to a resource Q column, and protein complexes were isolated by a 0-500mM NaCl gradient.
f. Analysis of eluted peak fractions presented in **(e)** by SDS-PAGE and siIver staining or immunoblotting using plakoglobin antibody. To identify the components that co-eluted with plakoglobin from the resource Q column, lane 7 in protein peak #5 was subjected to mass spectrometry analysis.
g. Distribution of proteins that co-eluted with plakoglobin from the anion exchange column (protein peak #5, lane 7 in **(e))** into DAVID-derived categories, and percentages of components assigned to each category.
h. lnteraction networks for components identified in **(g)** using the STRING database. Proteins are grouped by cellular distribution. Colors of individual proteins correspond to the distribution presented in **(g).** Active interactions (textmining, experiments, databases) are based on published data, using interaction score of high confidence (0.7).

To test this hypothesis, we loaded the eluates from the high MW protein peak on a Resource Q anion exchange column, and isolated protein complexes using a 0-500mM NaCl gradient. As shown in Fig. 1e and f, plakoglobin was eluted at a high yield together with several high MW polypeptides as a major monodisperse sharp peak at 250mM NaCl (peak #5). In addition to plakoglobin, our mass spectrometry analysis of this peak (Fig. 1f, lane 7) identified mainly cytoskeletal (28%) and membrane-bound (55%) proteins (Fig. 1g,h), including components comprising costameres, such as the DGC component delta-sarcoglycan, the basal lamina glycoprotein Laminin, which binds to the DGC, and the dystrophin associated protein spectrin^29^ (Fig. 1h and **Supplementary Table 1**). Furthermore, the major intermediate filament protein in muscle, desmin, was identified, as well as the associated proteins desmoplakin^30^, the desmin cross-linking protein plectin, and α-actinin, a component localized to the Z-bands where desmin filaments are aligned (Fig. 1h and **Supplementary Table 1**).

To confirm these findings by an independent approach, and identify additional proteins that bind plakoglobin, we transfected mouse Tibialis Anterior (TA) muscle with a plasmid encoding 6His-plakoglobin and isolated bound proteins with a Nickel column (Fig. 2a). Mass spectrometry analysis identified most of the aforementioned components as plakoglobin-bound proteins (**Supplementary Table 1**), and additional DGC core components, including dystrophin, α-sarcoglycan, β-sarcoglycan, γ-sarcoglycan, α-syntrophin, β-syntrophin, and β-dystroglycan, the costamere marker in skeletal muscle, vinculin, where DGC is normally located^13,15^, and the specialized lipid membrane domain caveolae markers, caveolin 1 and 3, which interact with DGC in striated muscle and link the cytoskeleton to the ECM^31,32^ (**Supplementary Table 2**). Other plasma membrane-bound proteins included insulin signaling components such as the p85 regulatory subunit of PI3K, which we previously showed can bind plakoglobin^23^, the insulin receptor substrate 1 (IRS1), which specifically binds to the insulin receptor and activates PI3K-Akt signaling^33^, two catalytic subunits of PI3K, type 3 and gamma, that bind p85-PI3K to form the PI3K complex, the insulin-like growth factor 2 receptor, and the insulin-like growth factor binding protein (**Supplementary Table 2**). These interactions of plakoglobin with components of the DGC and insulin signaling is consistent with its localization to the muscle membrane^23^ (Fig. 1, and see below), and with it playing an important role in striated muscle homeostasis^23,34,35^. Thus, in skeletal muscle plakoglobin seems to form multiprotein complexes containing structural and signal transmitting modules.

**Figure 2.**
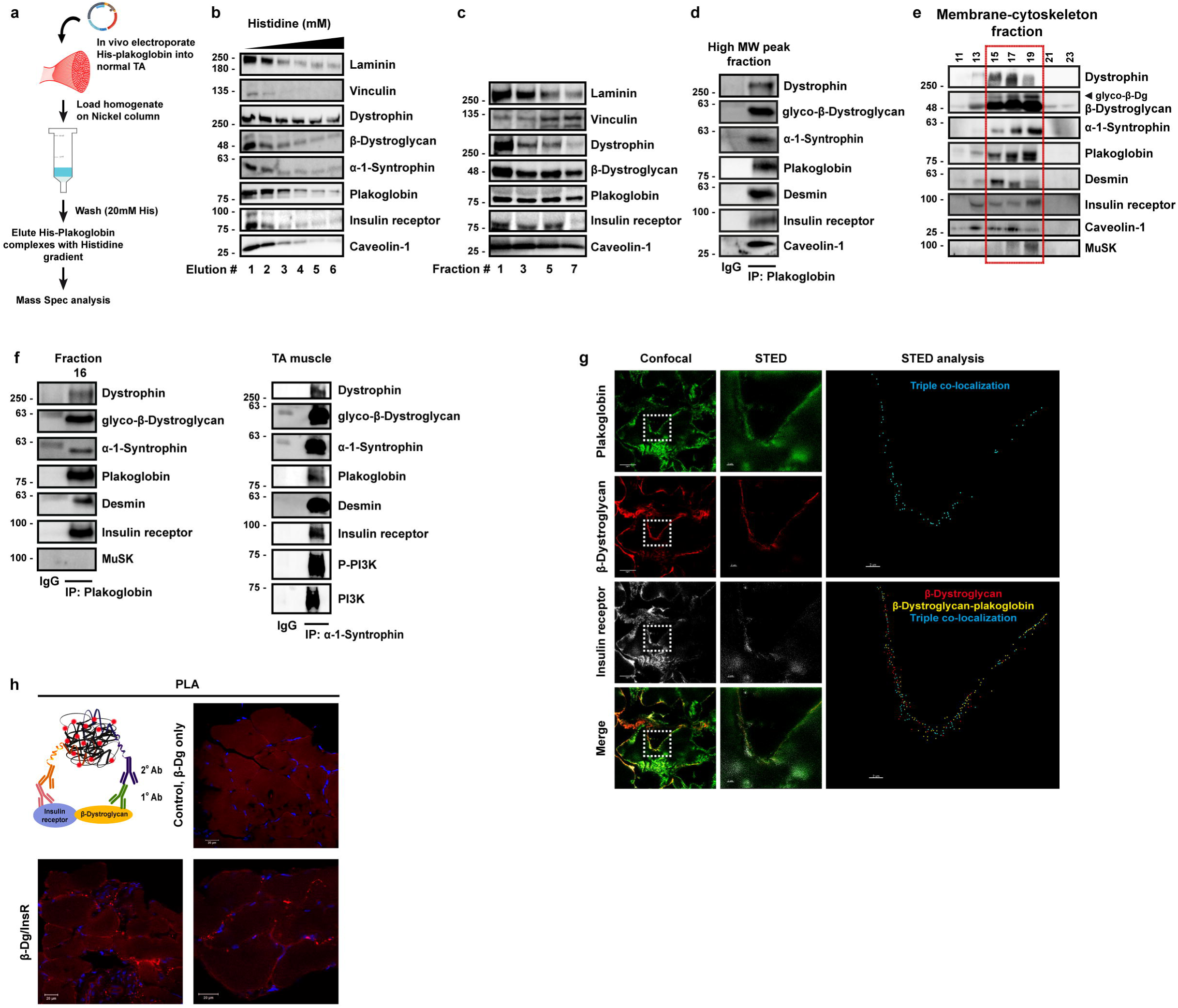
Plakoglobin binds DGC components and the insulin receptor at costameres on the plasma membrane. a. Affinity-based purification of plakoglobin-containing multiprotein assembly. A plasmid encoding 6His-tagged plakoglobin was electroporated into mouse TA mus cle, and 6His-plakoglobin-bound proteins were isolated with Nickel beads and identified by mass spectrometry.
b. His-plakoglobin-bound proteins were eluted from Nickel column in (a) with a Histidine gradient (50-250 mM) and analyzed by immunoblotting. Elution fraction #2 was subjected to mass spectrometry (one experiment was performed and data compared to the experiment presented in Figs. 1g,h).
c. Analysis of high MW protein peak fractions eluted from size exclusion chromatography (as in Fig. 1c) by SDS-PAGE and immunoblotting. n= two independent experiments.
d. Plakoglobin was immunoprecipitated from the high MW protein peak, and bound proteins were detected by immunoblotting.
e. Plakoglobin, DGC components (including glycosylated-β-dystroglycan), the insulin receptor, caveolin-1, desmin, and MuSK sediment to the same glycerol gradient fractions (marked bya red rectangle). Membrane-cytoskeleton fraction from mouse muscle was separated on 10-40% glycerol gradient, and alternate protein eluates were analyzed by Western blotting. n= two independent experiments.
f. In normal muscle, plakoglobin, DGC components, the insulin receptor, and desmin interact. Left: proteins co-purified with anti-plakoglobin from glycerol gradient fraction #16 illustrated in **(e)**, and detected by immunoblotting. MuSK did not bind plakoglobin although it sedimented to the same fractions in **(e).** Right: a reciprocal immunoprecipitation with a-1-syntrophin antibody from membrane fractions isolated from normal TA muscle. Two experiments were performed: one with plakoglobin antibody and one with a-1-syntrophin antibody.
g. Plakoglobin, β-dystroglycan, and the insulin receptor colocalize at costameres on skeletal muscle membrane. Confocal and STED images of TA muscle cross sections stained with the indicated antibodies. Scale bars, 5 µm (Confocal) and 2 µm (STED). n= three independent experiments indicating colocalization of these proteins. STED analysis is an annotated image in which all three proteins are detected using the spots module of lmaris software (white, red and green spheres). The double and triple co-occurrence spots are presented in yellow and blue, correspondingly.
h. Proximity ligation assay (PLA) was performed on TA cross sections with β-dystroglycan and insulin receptor antibodies or β-dystroglycan antibody alone. Red fluorescent dots indicate areas of β-dystroglycan-insulin receptor association.

### DGC, insulin receptors and plakoglobin form an intact multisubunit assembly

To learn whether DGC, insulin receptors and plakoglobin form one intact co-assembly, we repeated 6His-plakoglobin isolation from transfected TA muscles using a Nickel column, and eluted the bound proteins with a Histidine gradient. Western Blot analysis of the protein eluates showed that DGC components, vinculin, caveolin-1, the insulin receptor, and laminin were effectively co-purified with 6His-plakoglobin (Fig. 2b). These proteins physically interact *in vivo* at membrane caveolae because DGC components, insulin receptors, and caveolin-1 co-precipitated together with plakoglobin from the gel-filtration chromatography fractions of the high MW protein peak (Fig. 2c,d). These new interactions were corroborated by glycerol gradient fractionation of membrane-cytoskeleton preparations from TA muscle. Plakoglobin, DGC components, desmin, insulin receptor and p85-PI3K sedimented to the same fractions, as determined by the distribution of these proteins across the gradient (Fig. 2e). Interestingly, in these fractions β-dystroglycan appeared in its glycosylated form (Fig. 2e), which is essential for DGC stability^11^. These proteins interact within a native multimeric assembly because they co-precipitated together with plakoglobin from these glycerol gradient protein eluates, and also reciprocally with α-1-syntrophin (Fig. 2f). The presence of desmin in plakoglobin containing precipitates is consistent with the structural role of plakoglobin in linking the desmin cytoskeleton through membrane structures (e.g. DGC) to the ECM^36,37^. Moreover, these interactions were specific because Tyrosine Kinase Receptor, MuSK, which was present in the same glycerol gradient fractions together with DGC components, the insulin receptor, and plakoglobin (Fig. 2e), was not precipitated with plakoglobin antibody from muscle (Fig. 2f), heart or liver (see below).

We next determined if these novel interactions were unique to skeletal muscle or whether plakoglobin binds, and thus might regulate, these structural and signal transmitting modules generally. Surprisingly, similar interactions were demonstrated by coimmunoprecipitation in the heart and liver (**Supplementary Fig. 2**), and thus are probably playing important roles in many (perhaps all) tissues.

### The insulin receptor and DGC colocalize with plakoglobin on the sarcolemma

To determine whether plakoglobin, DGC and the insulin receptor colocalize on the plasma membrane, we performed immunofluorescence staining of muscle cross sections and analyzed by confocal and super-resolution stimulated emission depletion (STED) microscopy. Similar to epithelia^38^ and our previous observations in skeletal muscle^23^ plakoglobin showed a punctate distribution throughout the muscle fiber (Fig. 2g), and colocalized with β-dystroglycan and the insulin receptors (Fig. 2g and **Supplementary Fig. 3**), dystrophin and vinculin (**Supplementary Fig. 3a**), on the fiber membrane, which is consistent with the presence of these proteins in one intact protein assembly (Fig. 2b-f). Sub-diffraction spots colocalization analysis of STED images and three-dimensional analysis of confocal images confirmed a distinct heteromultimeric assembly of DGC-insulin receptor-plakoglobin with vinculin at costameres, which reside in the plasma membrane in register with the sarcomeric Z-bands (Fig. 2g and **Supplementary Fig. 3a,b**). These interactions were corroborated by a proximity ligation assay (PLA), which allows in situ detection of protein-protein interactions. Muscle cross sections were incubated with primary antibodies to insulin receptor and β-dystroglycan and secondary antibodies carrying complementary DNA probes. Because the targeted proteins are 20-100m apart, the probes could form a closed DNA circle, which served as a template for rolling circle amplification reaction, and subsequent incorporation of fluorescently labeled nucleotides (Fig. 2h). Consequently, the red fluorescent spots along the plane of the fiber membrane indicate areas where the insulin receptor and β-dystroglycan interact (Fig. 2h). These red spots were not formed in muscle sections incubated with β-dystroglycan antibody alone (Fig. 2h). Interestingly, these discrete regions where the insulin receptor, β-dystroglycan, and plakoglobin interact are in close proximity to the nucleus, most likely to efficiently translate signals from the plasma membrane (e.g. PI3K-Akt signaling) to transcriptional response (**Supplementary Fig. 3c**). Together, these findings indicate that in skeletal muscle, and probably other tissues, plakoglobin co-assembles with DGC and the insulin receptor on the plasma membrane, and may be important for bridging and stabilizing these structures.

**Figure 3.**
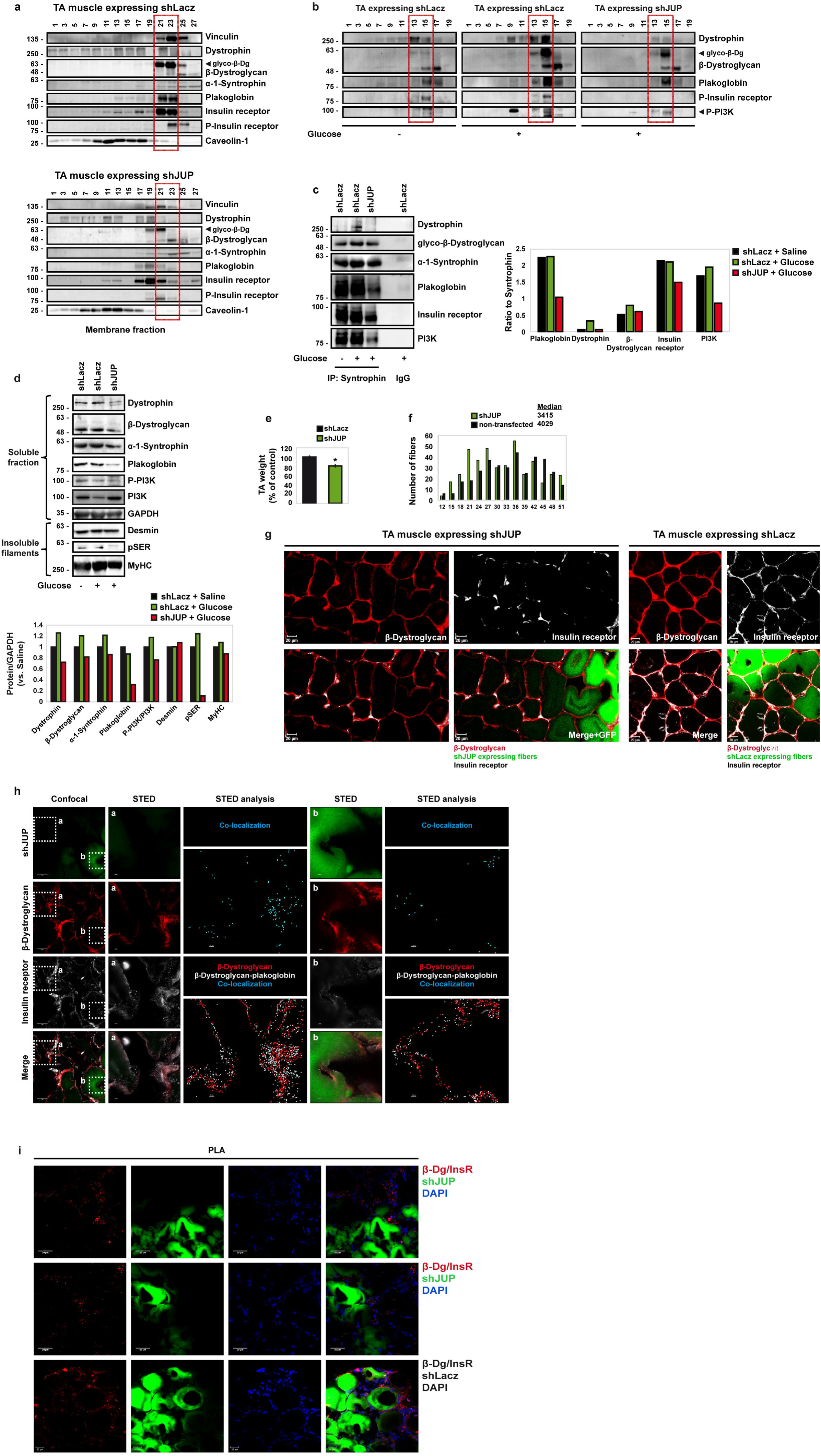
Plakoglobin downregulation reduces DGC-insulin receptor association and insulin signaling, and causes atrophy. a. Plakoglobin knockdown with plakoglobin shRNA (shJUP) promotes DGC-insulin receptor dissociation. TA mouse museles were transfected with shLacz or shJUP (contralateral limbs}, and purified membranes were separated on glycerol gradients and analyzed by SDS-PAGE and immunoblot. Red rectangle marks fractions that in shLacz expressing muscle contain plakoglobin complex with glycosylated-β-dystroglycan. n= three independent experiments.
b. Mice were injected with saline or glucose (1mg/gr body weight) and membrane fractions were isolated from transfected TA museles (contralateral limbs) and analyzed by glycerol gradient, and immunoblotting. Red rectangle marks fractions that in shLacz expressing musele contain plakoglobin complex with glycosylated-β-dystroglycan. n= three independent experiments.
c. Syntrophin was immunoprecipitated from purified membrane fractions of museles expressing shLacz or shJUP from mice injected with saline or glucose (1mg/gr body weight). Precipitates were analyzed by immunoblotting. Right: densitometric measurements of presented blots. Graph depicts ratio of each protein to syntrophin. n= two independent experiments.
d. Soluble fractions of TA museles expressing shLacz or shJUP from mice injected with saline or glucose were analyzed by SDS-PAGE and immunobloting. n= two independent experiments (representative blots are shown).
e. Downregulation of plakoglobin induces musele atrophy. Mean weights of TA museles expressing shJUP are plotted as the percentage of shLacz control museles. Data are represented as mean ± SEM. n = 14 mice; * P< 0.00005 by one-tailed t-test. n= two independent experiments.
f. Cross-sectional areas of 521 fibers transfected with shJUP (also express GFP, green bars) vs. 521 non-transfected fibers (black bars) in the same musele. n= 4 mice. Median is presented.
g. Representative confocal images of cross sections of museles expressing shLacz or shJUP (also express GFP}, stained with anti-insulin receptor and anti-β-dystroglycan. Scale bar, 20 µm. n= two independent experiments from two different mice.
h. Insulin receptor and β-dystroglycan colocalization is markedly reduced in muscle fibers expressing shJUP. Confocal and STED images of cross sections ofTA museles expressing shLacz or shJUP stained with the indicated antibodies. Scale bars, 5 µm (Confocal) and 2 µm (STED). n= two independent experiments. STED analysis is an annotated image in which the two proteins are detected using the spots module of lmaris software (white and red spheres). The double co-occurrence spots are presented in blue.
i. Insulin receptor and β-dystroglycan association is markedly reduced in musele fibers expressing shJUP. Proximity ligation assay (PLA) was performed on cross sections of TA museles expressing shLacz or shJUP stained with β-dystroglycan and insulin receptor antibodies. Red fluorescent dots indicate areas of β-dystroglycan-insulin receptor association.

### Insulin receptor activity is linked to DGC integrity via plakoglobin

To test whether plakoglobin enhances the structural integrity and functional properties of DGC and insulin receptor, we downregulated plakoglobin in normal muscle by electroporation of shRNA (shJUP)^23^, and analyzed the effects on DGC stability and insulin-PI3K-Akt signaling. Glycerol gradient fractionation of purified membranes from transfected muscles showed that most proteins sedimented to the same fractions as a one native complex, but not when plakoglobin was downregulated with shJUP (Fig. 3a). Plakoglobin loss led not only to decreased insulin receptor phosphorylation and activation, but also caused a marked reduction in glycosylated β-dystroglycan on the membrane indicating a pronounced destabilization of DGC^11^ (Fig. 3a). Furthermore, the content of vinculin, dystrophin, and the insulin receptor on the membrane was also reduced and their distribution was altered in shJUP expressing muscles strongly suggesting that the integrity of plakoglobin-containing complex was compromised (Fig. 3a). This reduction in the membrane content of DGC components and insulin receptor is selective and does not reflect a general reduction in total membrane protein because in the muscles expressing shJUP the protein levels of caveolin-1 and the unmodified form of β-dystroglycan were similar to shLacz controls (Fig. 3a).

Consistent with these results, glucose injection to mice to induce insulin secretion enhanced phosphorylation of the insulin receptor and PI3K in control muscles (compared with mice injected with saline)(Fig. 3b), and promoted association of dystrophin, glycosylated-β-dystroglycan, plakoglobin, insulin receptor and PI3K because more of these proteins sedimented to the same glycerol gradient fractions (Fig. 3b) and they could be co-precipitated with the α-1-syntrophin (Fig. 3c). This association required plakoglobin because in the muscles expressing shJUP, insulin receptor and PI3K phosphorylation was markedly reduced, and much less of these proteins were associated in one intact complex (Fig. 3b,c). Similar effects were obtained in muscles expressing shJUP or shLacz from mice injected with saline (**Supplementary Fig. 4a**). Interestingly, the soluble pools (the fraction soluble at 6,000 g) of DGC subunits (i.e. dystrophin, β-dystroglycan, α-1-syntrophin) increased on glucose injection but were smaller when plakoglobin was downregulated and insulin signaling fell (Fig. 3a,b,d), probably due to their degradation by the proteasome (Fig. 3d). Moreover, loss of plakoglobin and the resulting reduction in DGC integrity and its association with the insulin receptor enhanced depolymerization of the desmin cytoskeleton because the amount of phosphorylated desmin filaments decreased in the insoluble fraction (Fig. 3d)^39,40^. In fact, these effects by shJUP must underestimate the actual effects of plakoglobin downregulation because only about 70% of the fibers were transfected (**Supplementary Fig. 4b**). Thus, in normal muscle, plakoglobin is required for DGC stability, which appears to be linked to normal signaling through the insulin receptor.

**Figure 4.**
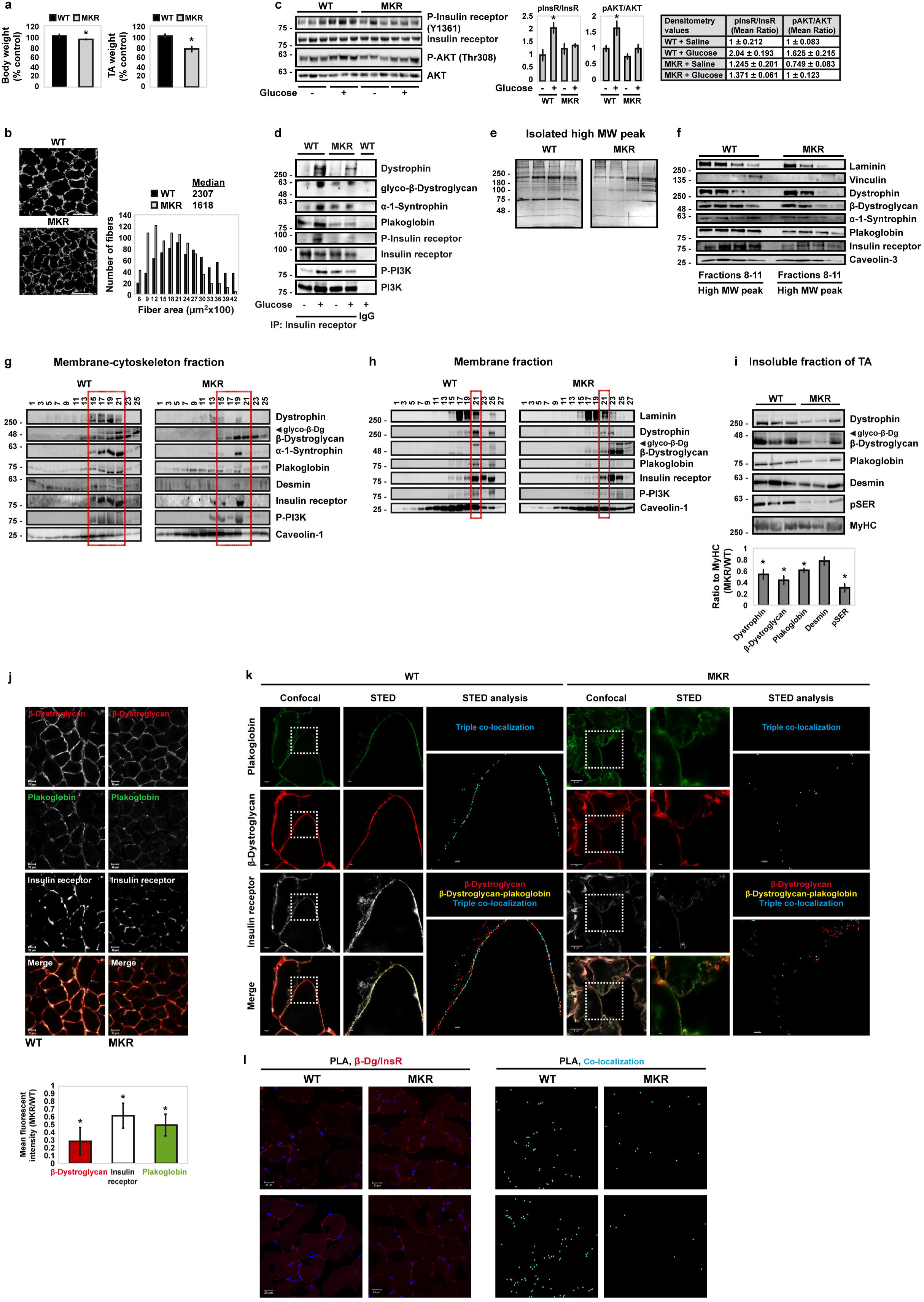
In museles from MKR mice insulin receptor-DGC association is markedly reduced. a. Mean body and TA weights of MKR mice at 8 weeks of age are presented as percentage of WT littermates. Body weights, n = 5 mice; TA weights, n = 12 mice; * P < 0.05, by one-tailed t-test. Data are represented as mean ± SEM. n= two independent experiments.
b. Cross-sections of TA museles from MKR mice are smaller than WT. Left: laminin staining in white. Scale bar, 50 µm. Right: Cross-sectional areas of 834 fibers from WT (black bars) and MKR (grey bars) mice are presented. n= 3 mice. Median is presented.
c. Insulin signaling in MKR mice museles is abolished. Left: soluble fractions of museles from WT and diabetic MKR mice injected with saline or glucose (1mg/gr body weight) were analyzed by SDS-PAGE and immunoblot. n= two independent experiments. Each lane represents one musele from one mouse. Right: densitometric measurement of presented blots is depicted by graph and densitometry values.
d. In MKR mice musele, DGC-insulin receptor association is reduced. Insulin receptor immunoprecipitation from membrane fractions of WT and MKR mice museles injected with saline or glucose (1mg/gr body weight). n= two independent experiments.
e. High MW protein peak fractions isolated by size-exelusion chromatography from WT and MKR mice museles were analyzed by SDS-PAGE and silver staining in three independent experiments (a representative is shown).
f. Fractions in **(e)** were analyzed by immunoblotting.
g. Membrane-cytoskeleton fractions from WT and MKR mice TA museles were analyzed by glycerol gradient fractionation. Red rectangle marks fractions that in WT mouse musele contain plakoglobin complex with glycosylated-β-dystroglycan. n= two independent experiments.
h. Isolated membranes from WT and MKR mice museles were analyzed by glycerol gradient fractionation. n= two independent experiments. Red rectangle marks fractions that in WT mouse musele contain plakoglobin complex with glycosylated-β-dystroglycan.
i. Top: insoluble fractions of TA museles from WT and MKR mice were analyzed by immunoblotting in two independent experiments (a representative is shown). Each lane represents one musele from one mouse. Bottom: densitometric measurement of presented blots.
j. Top: Cross sections of WT and MKR mice TA museles were stained with plakoglobin, β-dystroglycan and insulin receptor antibodies. Scale bar, 20 µm. n= three independent experiments (representative confocal images are shown). Bottom: fluorescent intensity was quantified using Imaris software, and data from three independent experiments in depicted in a graph.
k. Plakoglobin, insulin receptor, and β-dystroglycan colocalization is markedly reduced in museles from MKR mice. Confocal and STED images of cross sections of TA museles from WT and MKR mice stained with the indicated antibodies. Scale bars, 5 µm (Confocal) and 2 µm (STED). n= two independent experiments. STED analysis is an annotated image in which all three proteins are detected using the spots module of Imaris software (white, red and green spheres). The double and triple co-occurrence spots are presented in yellow and blue, correspondingly.
l. Insulin receptor and β-dystroglycan association is markedly reduced in MKR mice museles. Proximity ligation assay (PLA) was performed on cross sections of TA museles from WT or MKR mice stained with β-dystroglycan and insulin receptor antibodies. Red fluorescent dots (left) and blue dot analysis (right) indicate areas of β-dystroglycan-insulin receptor association.

As expected from the fall in PI3K-Akt signaling, in the muscles expressing shJUP atrophy became evident because the mean TA weights was lower by 20% than control (Fig. 3e), and the mean cross-sectional area of 500 fibers expressing shJUP was smaller than 500 non-transfected ones (Fig. 3f), as reported^23^. Finally, immunofluorescence staining using confocal (Fig. 3g) or super-resolution STED (Fig. 3h) microscopy, and PLA analysis (Fig. 3i) of muscles expressing shJUP or shLacz confirmed colocalization and association of the insulin receptor and β-dystroglycan on the sarcolemma of non-transfected and shLacz expressing fibers but significantly less in those muscle fibers expressing shJUP. In the shJUP fibers, the staining intensity of insulin receptor and its association with β-dystroglycan were reduced compared with non-transfected fibers (Fig. 3g,h,i), while only a slight reduction in β-dystroglycan could be observed because, as demonstrated by our biochemical analyses (Fig. 3a,b and **Supplementary Fig. 4a**), plakoglobin downregulation markedly reduced the amount of glycosylated-β-dystroglycan on the plasma membrane but the unmodified form of β-dystroglycan was quite similar to shLacz controls. In the shLacz expressing fibers, however, the staining intensities of insulin receptor and β-dystroglycan and their association were similar to the adjacent non-transfected fibers (Fig. 3g,i). Thus, plakoglobin is required for DGC stability and association with insulin receptors, and for normal PI3K-Akt activity.

### Perturbation of insulin receptor activity reduces its association with the DGC and plakoglobin

Altogether, these observations suggest that DGC integrity is linked to insulin receptor activity via plakoglobin. To investigate whether the content of DGC on the membrane is influenced by the fall in PI3K-Akt signaling, we used transgenic mice (MKR) model for type-2 diabetes, which express a kinase inactive form of insulin-like growth factor 1 receptor (IGF-1R) specifically in skeletal muscle. This IGF-1R mutant dimerizes and inhibits IGF-1R and insulin receptor activities ultimately leading to skeletal muscle-specific inhibition of PI3K-Akt signaling, and insulin resistance^4,41^. Consistent with prior studies^41^, at 8 weeks of age, the mean body weights of MKR mice were 7% lower, the mean wet weights of their TA muscles were 25% lower, and the cross-sectional area of their muscle fibers was smaller than age-matched control mice (Fig. 4a,b). In MKR mice muscles PI3K-Akt signaling falls^41^. Accordingly, glucose-stimulated insulin secretion induced phosphorylation of the insulin receptor and Akt, and increased coprecipitation of PI3K, plakoglobin, and DGC components with the insulin receptor in muscles from WT mice (Fig. 4c,d). However, in muscles from MKR mice, these interactions were markedly reduced and insulin signaling was not activated upon glucose injection (Fig. 4c,d). Thus, impaired insulin receptor function reduces its association with the DGC.

To learn how this fall in PI3K-Akt signaling influences DGC integrity, we analyzed muscles from MKR and WT mice using gel filtration chromatography (as in Fig. 1). Similar to our observations above, the high MW protein peak fractions from normal muscle homogenates contained mostly components of a high MW, which comprised the large multiprotein plakoglobin-containing complex (Fig. 4e,f). However, in MKR mice muscles, this protein peak contained lower amounts of plakoglobin, DGC components, and insulin receptors than in WT littermates (Fig. 4e,f), suggesting that MKR mice muscles the amount of the plakoglobin-containing complex is reduced. Although caveolin-3 levels in MKR muscles were similar to WT, the amounts of laminin and vinculin were reduced suggesting that in MKR mice muscles the structural link between the cytoskeleton and laminin at the ECM, via vinculin and DGC, is perturbed (Fig. 4f). The proteins whose amount was reduced in the cytoskeleton-membrane fraction of muscles from MKR mice are probably degraded because they did not accumulate in the soluble fraction (**Supplementary Fig. 5a**).

**Figure 5.**
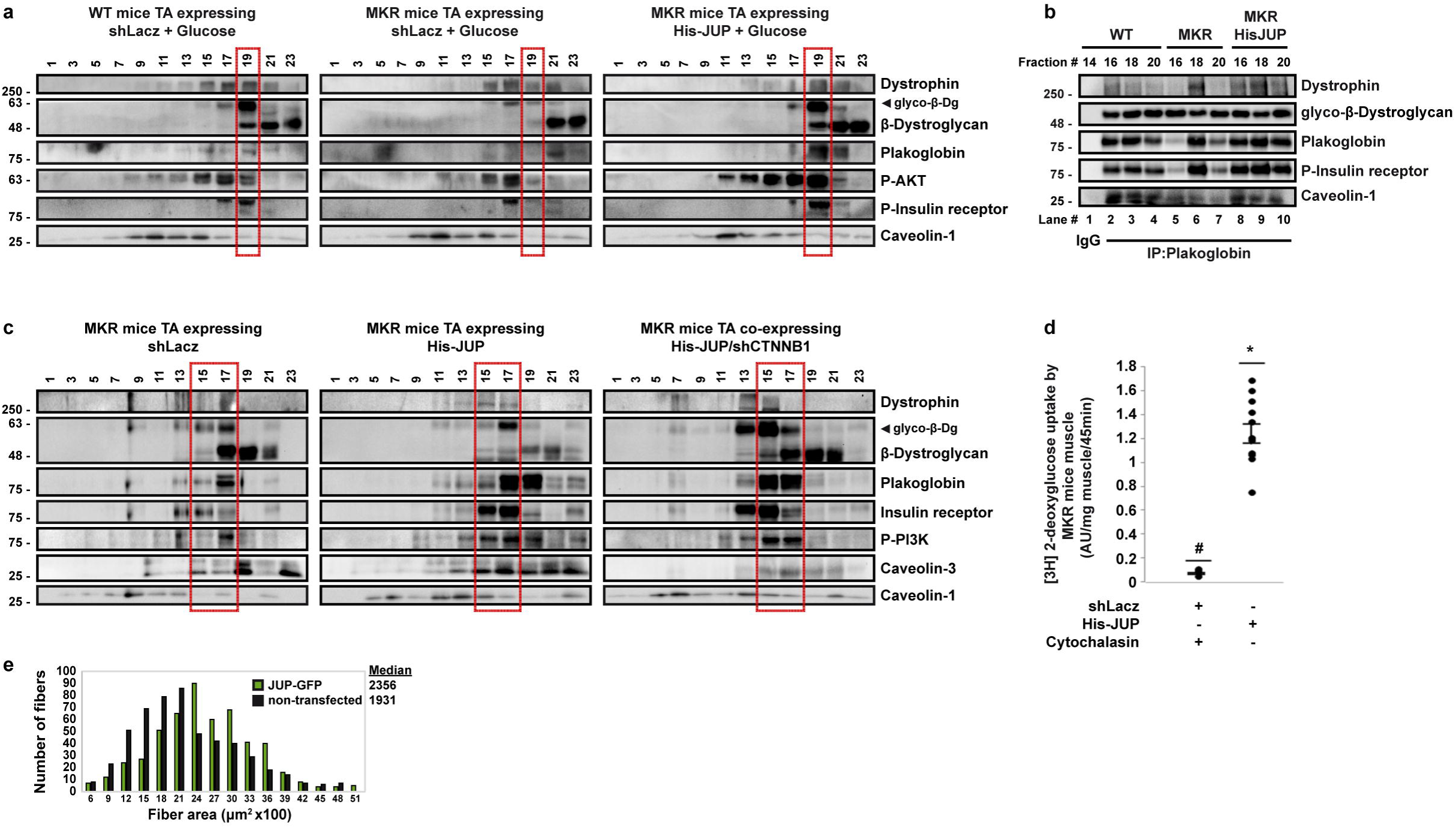
Overexpression of plakoglobin alone enhances DGC-insulin receptor association and insulin signaling in museles from MKR mice. a. Mice were injected with glucose (1mg/gr body weight) and purified membrane fractions from WT and diabetic MKR mice museles expressing shLacz or 6His-plakoglobin (His-JUP) were analyzed by glycerol gradient, SDS-PAGE and immunoblot. In MKR mice, contralateral limbs were transfected. n= three independent experiments.
b. Plakoglobin overexpression in MKR mice museles enhances DGC-insulin receptor association and insulin receptor activity. Plakoglobin immunoprecipitation from glycerol gradient fractions presented in **(a)** was analyzed by immunoblotting. n= two independent experiments.
c. The beneficial effect of plakoglobin on DGC-insulin receptor association does not require its close homolog β-catenin. Membrane fractions purified from MKR mice museles expressing shLacz, His-plakoglobin, or His-plakoglobin together with β-catenin shRNA (shCTNNB1) were analyzed by glycerol gradient fractionation and immunoblotting. MKR mice museles transfected with His-plakoglobin, or His-plakoglobin together with shCTNNB1 are from contralateral limbs of one mouse. n= two independent experiments.
d. Overexpression of plakoglobin in museles from diabetic MKR mice enhances glucose uptake. [3H]-2-deoxyglucose uptake by MKR mice museles expressing shLacz in the presence of Cytochalasin B (a competitive inhibitor of regulated glucose transport into cells) or 6His-plakoglobin (His-JUP) is plotted as ratio to shLacz control, which was not treated with Cytochalasin B. Data presented as AU/mg muscle/45 min. Data are mean ± SEM, n=11 mice for His-JUP and shl acz, n=4 mice for shLacz treated with Cytochalasin B, • p=0.01 and # p=0 vs. shLacz in the absence of Cytochalasin B, by one-tailed t-test. n= two independent experiments.
e. Overexpression of plakoglobin in museles from MKR mice attenuates the reduction in fiber size. Measurement of cross-sectional area of 530 fibers transfected with His-JUP (and expressing GFP; green bars) vs. 530 nontransfected fibers (black bars) in the same muscle. Data acquired from seven mice. Median is presented.

This reduced content of the DGC in muscles from MKR mice where insulin signaling is impaired was corroborated biochemically by glycerol gradient fractionation of membrane-cytoskeleton preparations (Fig. 4g) and purified membranes (Fig. 4h). In muscles from WT mice, DGC components, plakoglobin, the insulin receptor, PI3K and desmin sedimented to the same fractions (Fig. 4g,h) suggesting that they formed an intact complex (Fig. 2d,f) in which plakoglobin functions as a molecular linker (Fig. 3c). Conversely, the amount of this multi-protein assembly was markedly reduced in muscles from MKR mice compared with WT probably because they are degraded in those muscles lacking active insulin receptor (Fig. 4g,h). In addition, the amount of glycosylated β-dystroglycan decreased in MKR mice muscles, and instead β-dystroglycan accumulated in its non-glycosylated form (Fig. 4g,h). This reduction in glycosylation and DGC amount in muscles where PI3K-Akt signaling is impaired, is most probably due to the fall in plakoglobin content on the membrane (Fig. 4d,h), which is sufficient to cause DGC-insulin receptor dissociation (Fig. 3). Thus, plakoglobin loss or reduced function^23^ may be an early event leading to impaired insulin signaling and loss of intact DGC in atrophy or disease.

In the insoluble fraction (6000 g pellet) of muscles from MKR mice, phosphorylated desmin filaments also decreased in amount compared with control (Fig. 4i), suggesting that this cytoskeletal network depolymerize under conditions where DGC content is low and insulin signaling falls. This reduction in phosphorylated species of desmin filaments indicates enhanced depolymerization (Fig. 4i), as occurs in atrophying muscles from mouse and human^39,40,42,43^, and not reduced gene expression because the mRNA levels of desmin were similar in muscles from WT and MKR mice (**Supplementary Fig. 5b**). Furthermore, components comprising plakoglobin-containing complex bound to this insoluble pellet in muscles from WT mice (Fig. 4i), consistent with the structural role of the desmin cytoskeleton in linking the myofibrils to membrane structures (e.g. DGC). However, the insoluble pellets from MKR mice muscles contained fewer components comprising this multi-subunit complex and their loss exceeded the reduction in myofibrillar myosin heavy chain (MyHC)(Fig. 4i), which should be reflected by an overall reduction in the number of these protein clusters on the muscle membrane. In fact, immunofluorescence staining of TA cross-sections and analyses by Confocal (Fig. 4j) and super-resolution STED (Fig. 4k) microscopy indicated a colocalization of plakoglobin with, insulin receptor, β-dystroglycan, and vinculin on the plane of the fiber membrane, which was markedly reduced in MKR mice, where plakoglobin seemed to accumulate inside the muscle fiber (Fig. 4j,k, and **Supplementary Fig. 5c,d**). This reduction in the mean fluorescent intensity of plakoglobin, insulin receptor, and β-dystroglycan is selective because the average fluorescent intensity of Wheat Germ Agglutinin (WGA) in the same muscles was not lower than in WT controls (**Supplementary Fig. 5e**). In addition, sub-diffraction spots colocalization analysis of STED images indicated a marked decrease in the number of protein clusters (Fig. 4k), and PLA analysis using insulin receptor and β-dystroglycan antibodies indicated a significant reduction in insulin receptor-β-dystroglycan association (Fig. 4l) in muscles from MKR mice compared with WT. Thus, the impaired functions of the insulin and IGF-1 receptors in MKR mice muscles, and the resulting fall in PI3K-Akt signaling ultimately lead to a reduction in DGC content, DGC-insulin receptor clusters, and muscle fiber size.

### Plakoglobin overexpression in muscles from MKR mice enhances DGC-insulin receptor association and PI3K-Akt activity

Together, these observations suggested that increasing the level of plakoglobin in MKR mice muscle should enhance DGC-insulin receptor association, accentuate PI3K-Akt activity, and thus increase sensitivity to insulin. Accordingly, plakoglobin-DGC-insulin receptor heteromultimers were compromised and PI3K-Akt signaling was low in muscles from MKR mice compared with WT (Fig. 5a), but not when 6His-plakoglobin was overexpressed (Fig. 5a). Plakoglobin accumulation not only enhanced insulin-stimulated PI3K-Akt signaling, but also caused a greater interaction between plakoglobin, DGC components, the insulin receptor and p85-PI3K in MKR mice muscles than in control (Fig. 5b). Interestingly, although insulin receptor-plakoglobin association was markedly reduced in MKR mice muscles, β-dystroglycan remained bound to plakoglobin (Fig. 5b, lanes 5-7) and their association was even greater when plakoglobin levels were low (lanes 5 and 7, ratio of β-dystroglycan to plakoglobin), suggesting an altered distribution of β-dystroglycan on the membrane when its association with the insulin receptor and other DGC components (e.g. dystrophin) is low (Fig. 5b). It is noteworthy that this beneficial effect of plakoglobin overexpression on DGC-insulin receptor association and insulin signaling did not require the function of plakoglobin close homolog, β-catenin, because in the MKR mice muscles expressing 6His-tagged plakoglobin, transfection of β-catenin shRNA (shCTNNB1) to simultaneously downregulate β-catenin did not perturb plakoglobin-DGC-insulin receptor association (Fig. 5c). Thus, plakoglobin is a novel auxiliary subunit linking insulin receptor activity to DGC integrity. These beneficial effects on insulin signaling by plakoglobin overexpression also enhanced muscle glucose uptake in MKR mice (Fig. 5d)(Cytochalasin B is a cell-permeable mycotoxin that competitively inhibits glucose transport into cells, and was used here to evaluate specifically regulated glucose uptake^44^), and markedly attenuated muscle fiber atrophy (Fig. 5e). Thus, plakoglobin mediates insulin-dependent glucose uptake and regulates muscle size by stabilizing insulin receptor-DGC association.

### DGC, insulin receptors and plakoglobin interact through the armadillo repeats and the N-terminal domain of plakoglobin

Co-assembly of DGC, the insulin receptor and plakoglobin was finally demonstrated by mapping the sites that constitute their interaction. Plakoglobin is composed of an N-terminal head domain, a C-terminal tail domain, and a central conserved domain containing 12 armadillo repeats, each of 42 amino acids in length^45^. Mutations within the N-terminal domain reduce plakoglobin membrane localization and cause cardiomyopathy in human^34^, and deletion of the C-terminal domain leads to the formation of large abnormal cadherin-based membrane structures and causes Naxos disease^46,47^. To identify the critical domains within plakoglobin, which are required for its heterodimerization with DGC and the insulin receptor, we generated 6His-V5-tagged truncation mutants of plakoglobin lacking the N-terminal domain (ΔN), the C-terminal domains (ΔC), or either end domain (ΔNC), while leaving the central armadillo domain intact (Fig. 6a). Overexpression of ΔN-plakoglobin in normal mouse TA for 6 d resulted in decreased insulin receptor phosphorylation compared with muscles expressing full-length (FL) plakoglobin, strongly suggesting that the N-terminus of plakoglobin is required for its interaction with and activation of the insulin receptor (Fig. 6b). Affinity purification of plakoglobin truncation mutants from transfected muscles using Nickel column and histidine gradient (Fig. 6c), and subsequent densitometric measurement of protein ratios (Fig. 6d) indicated that deletion of plakoglobin N-terminus (ΔN) reduced its association only with the insulin receptor, but not with β-dystroglycan or dystrophin, compared with FL-plakoglobin (Fig. 6d), which is consistent with the reduction in insulin receptor activity (Fig. 6b). By contrast, loss of plakoglobin C-terminus (ΔC) has been reported to promote the formation of large abnormal membrane structures in Naxos disease^46,47^, and accordingly enhanced association of this mutant with both the insulin receptor and β-dystroglycan was observed (Fig. 6d). In addition, the deletion of plakoglobin C-terminus in ΔNC mutant seemed to compensate for the loss of plakoglobin N-terminus because this mutant remained bound to the insulin receptor (Fig. 6d). Plakoglobin C-terminus probably masks protein interaction sites that are close to its N-terminus, as was proposed in cardiac muscle^46,48^. These interactions were specific because none of these proteins bound the Nickel column in control muscles expressing shLacz (Fig. 6d). Therefore, insulin receptor interacts with plakoglobin N-terminus, and β-dystroglycan binds to sites adjacent to this region, which seem to be regulated by plakoglobin C-terminal tail. Interestingly, dystrophin bound equally to FL-plakoglobin and all truncation mutants (Fig. 6d) suggesting that its binding sites are located within plakoglobin central armadillo repeats domain. Thus, plakoglobin appears to function as a critical scaffold protein linking DGC and insulin receptors physically and functionally to maintain normal muscle mass.

**Figure 6.**
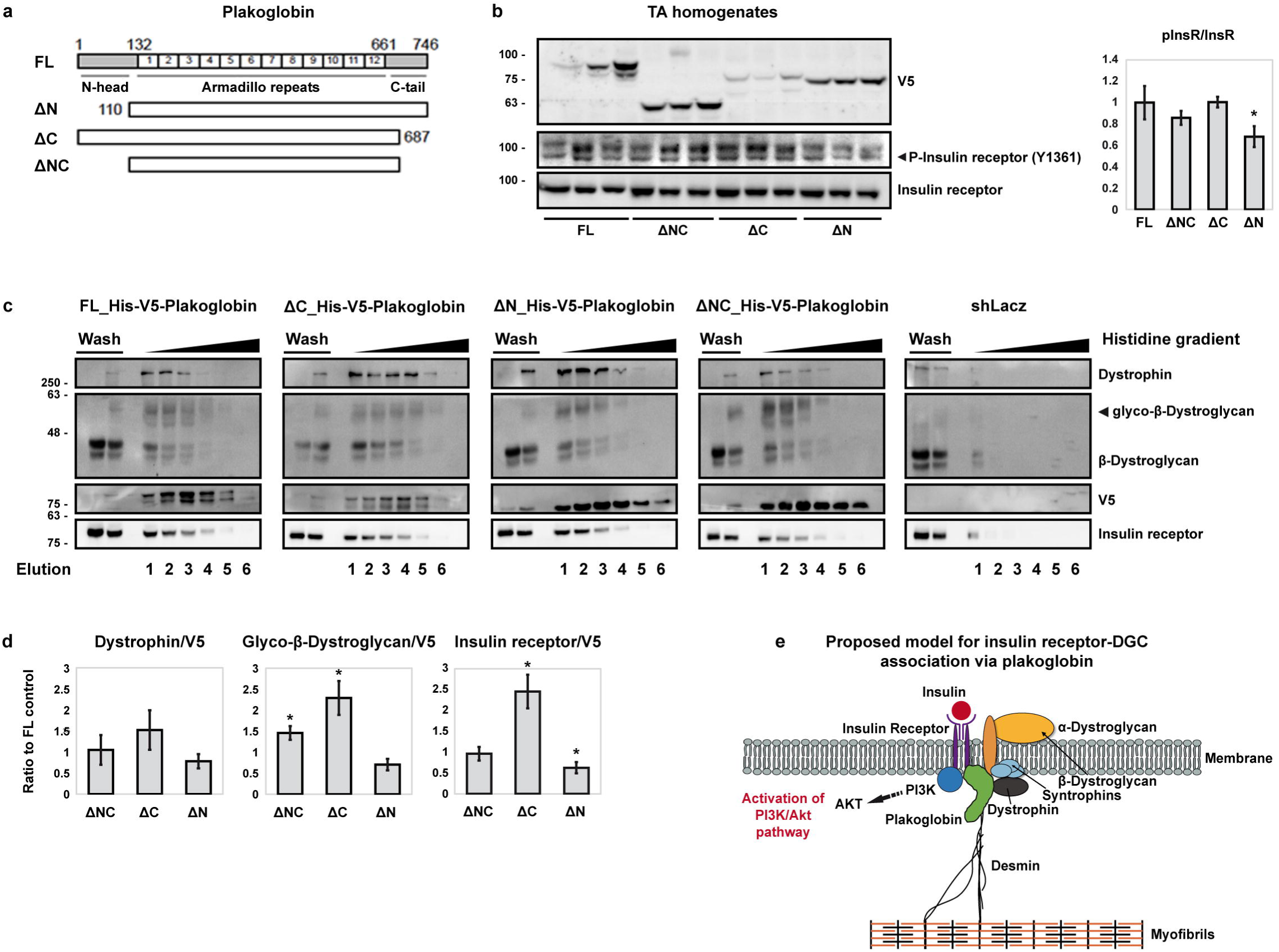
ldentification of interaction sites between plakoglobin, β-dystroglycan and the insulin receptor. a. Illustration of plakoglobin truncation mutants. **(b,c,d)** Insulin receptor binds to plakoglobin N terminus, and adjacent regions in plakoglobin are recognized by β-dystroglycan. TA museles from fed mice were electroporated with plasmids encoding 6His-V5-plakoglobin (FL), or its truncation mutants lacking the C-terminus (ΔC), the N-terminus (ΔN) or both (ΔNC).
b. Loss of plakoglobin N-terminus reduces insulin receptor activity. Left: soluble extracts from transfected museles were analyzed by immunoblotting using specific antibodies. Each lane represents one muscle from one mouse. n= two independent experiments (a representative blot is shown). Right: densitometric measurement of presented blots. Data are mean ± SEM, n=3. * p<0.05. vs. FL, by one-tailed t-test.
c. Isolated membrane fractions from transfected TA museles were loaded on Nickel column in parallel, and bound proteins were purified with histidine gradient. Lanes 1-6 represent elution fractions # 1-6 with increasing histidine concentration (20-250mM). Data is compared to control muscle expressing shLacz. n= five independent experiments (representative blots are shown).
d. Densitometric measurement of insulin receptor, glycosylated-β-dystroglycan, and dystrophin ratios to V5-tagged plakoglobin mutants in the Nickel column eluates. Ratios to V5-tagged FL-plakoglobin are presented. Data are mean ± SEM, n=S. * p < 0. 05. vs. FL, by one-tailed t-test.
e. Proposed model for DGC-insulin receptor association via plakoglobin.

## DISCUSSION

These studies have uncovered novel protein assemblies of insulin receptors and DGC, whose structural integrity and functional properties are linked and endowed by plakoglobin. We show that plakoglobin stabilizes these structural and signaling modules at costameres, and contributes to maintenance of cell size by promoting tissue integrity through desmin-DGC-laminin axis, and simultaneously regulating signaling through the insulin receptor. Plakoglobin’s stimulation of insulin signaling seems to result from it promoting a tight assembly between the insulin receptor and DGC because decreasing plakoglobin levels in normal muscle reduced DGC-insulin receptor association and insulin signaling, whereas increasing plakoglobin content in diabetic mice muscles enhanced DGC-insulin receptor co-assembly and glucose uptake. Consequently, DGC acts as a cellular signaling node containing plakoglobin as a pivotal subunit, and perturbation of plakoglobin function (e.g. by Trim32^23^) probably contributes to the insulin resistance in DMD^7,8^ or metabolic disorders (e.g. untreated diabetes, obesity).

Plakoglobin has been originally described as a component of desmosomes that are prominent in tissues that must withstand mechanical stress (e.g. heart), but its role in skeletal muscle had been generally overlooked. Stimulation of insulin secretion by the injection of glucose enhanced association of dystrophin and glycosylated-β-dystroglycan with 1-α-Syntrophin, attracted PI3K to this complex, and activated insulin signaling, but not when plakoglobin was downregulated with a specific shRNA (Fig. 3b,c). The glycosylation of β-dystroglycan, which is essential for DGC stability^11^, seems to require plakoglobin because in those shJUP expressing muscles the levels of glycosylated-β-dystroglycan were markedly reduced (Fig. 3a,b,c and **Supplementary Fig. 4a**). Under normal conditions, when growth or survival signals are transmitted, plakoglobin may promote trafficking of glycosylated-β-dystroglycan from the golgi apparatus to the plasma membrane, or prevent its recycling from the plasma membrane, in order to maintain DGC stability. Under stress conditions, however, reduced amount of plakoglobin on the plasma membrane (Fig. 4) may enhance loss of glycosylated-β-dystroglycan and DGC recycling or alternatively reduce the trafficking of DGC components to and their assembly on the plasma membrane. Consistently, in MKR mice muscles, impaired insulin receptor activity led to a prominent reduction in the amounts of plakoglobin and DGC components on the membrane (Fig. 4), while these proteins did not accumulate in the cytosol (**Supplementary Fig. 5a**). This marked loss occured presumably before the trafficking of these components from golgi to the plasma membrane or after they dissociated from each other in muscles lacking active insulin receptor. Therefore, a reciprocal regulation of DGC and the insulin receptor seems to be required to promote DGC stability and insulin receptor activity. β-Dystroglycan may function as a scaffold for the insulin receptor as it does for Acetylcholine Receptor during the formation of Neuromuscular Junctions in development^49^, or as a signaling hub maintaining insulin receptor activity and expression (by affecting gene expression as suggested for other genes^50^). In turn, active insulin receptor may stabilize the DGC by phosphorylation of its components^50^ or by inhibiting their degradation via the proteasome or autophagy (by activating insulin-PI3K-Akt-FoxO pathway^51^). Alternatively, PI3K-Akt signaling may activate transcription factors that increase gene expression of DGC components, and the newly synthesized proteins serve as precursors for DGC assembly on the plasma membrane. Thus, impaired insulin receptor activity (e.g. in MKR mice muscles) may result in reduced expression or increased proteolysis of DGC components and the insulin receptor itself, and consequently reduced amount of DGC-insulin receptor assemblies on the plasma membrane.

The beneficial effects of plakoglobin overexpression on DGC-insulin receptor association seem to be more pronounced in the muscle co-transfected with β-catenin shRNA than in muscles expressing His-plakoglobin alone (Fig. 5c). These effects most likely result from the higher transfection efficiency of His-plakoglobin in the co-transfected muscles rather than from the simultaneous downregulation of β-catenin. However, it is also possible that plakoglobin functions in part through regulation of gene expression (as proposed^52,53^), and competes with its homolog β-catenin for binding to the LEF/TCF family of transcription factors. In this case, plakoglobin overexpression with a simultaneous downregulation of β-catenin might enhance the transcriptional activity of plakoglobin. The physiological significance of plakoglobin as a regulator of gene expression remains controversial, and its potential transcriptional role in skeletal muscle and the effects on DGC-insulin receptor association are important questions that are beyond the scope of this study. In any case, the stabilization of DGC-insulin receptor association by plakoglobin overexpression suggests that increasing plakoglobin content can surpass certain inhibitory effects imposed on this protein to prevent its association with DGC and insulin receptors. The resulting increase in insulin signaling will accentuate the cellular response to insulin and thus enhance protein synthesis and glucose uptake, reduce protein degradation, and promote growth. Because plakoglobin is expressed in all cells, and since similar interactions were demonstrated in heart and liver (**Supplementary Fig. 2**), these functions of plakoglobin are probably general to many tissues.

This study shows for the first time that cellular metabolism is tightly linked to tissue structural properties via plakoglobin. Plakoglobin serves as scaffold protein at the sarcolemma, where the insulin receptor is positioned to interact with the DGC. Our investigation of binding sites on plakoglobin revealed that insulin receptor interacts with plakoglobin N-terminus and β-dystroglycan recognizes adjacent domains, which seem to be masked by plakoglobin C-terminal region^46,48^. Deletion of plakoglobin’s C-terminal domain causes Naxos disease and leads to the formation of large abnormal cadherin-based membrane structures, suggesting that this domain limits the size of membrane structures and regulates protein-protein interactions required for their assembly^46,47^. Consistently, we demonstrate that overexpression of plakoglobin ΔC mutant enhances association with the DGC and the insulin receptor on the plasma membrane, albeit with no significant effects on insulin receptor activity (Fig. 6). In knock-in mice expressing the C-terminus deletion mutant of plakoglobin cardiac function was normal suggesting that this carboxy-terminal domain is dispensable for plakoglobin function in the heart^54^. Thus, the effects of this mutation on plakoglobin function and muscle architecture and its role in cardiomyopathies remains debated and require in-depth study based upon the present identification of the novel complexes that plakoglobin forms in skeletal and cardiac muscles, and its physiological roles.

In the heart and epithelia, plakoglobin links desmin filaments to desmosome complexes via desmoplakin^36,37,55^. Although skeletal muscle lacks “classic” desmosomes, it clearly contains desmosomal components, which are present on the surface membrane and can be isolated together with plakoglobin, DGC and insulin signaling components (**Supplementary Table 1**)^23^. Unlike β-dystroglycan, plakoglobin is not evenly distributed along the sarcolemma, and instead accumulates in discrete regions by binding to specific receptors to stimulate signaling pathways and to promote cell–ECM contacts as proposed here. These regions are found in close proximity to the nucleus probably to efficiently translate extracellular signals to transcriptional response. Furthermore, these clusters of plakoglobin-DGC-insulin receptor contain vinculin, which in skeletal muscle is confined to costameres that link the desmin cytoskeleton and the bound contractile apparatus to laminin at the ECM thus contributing to the mechanical integrity of the muscle tissue^16^. In fact, the strongest evidence for the importance of plakoglobin to desmin filament stability and thus tissue architecture were our findings that in normal muscle plakoglobin downregulation alone was sufficient to promote desmin depolymerization and muscle atrophy (Fig. 3d,e,f). Whether loss of intact desmin filaments in skeletal muscle, as occurs in desmin myopathies, affects DGC integrity or signal transmission via the insulin receptor, and whether plakoglobin cellular distribution changes in these diseases, require further study.

Plakoglobin may mediate growth promoting signals also via the IGF1-R (**Supplementary Table 1**), although plakoglobin’s beneficial effects on insulin signaling seem to result primarily from it acting on the insulin receptor because in MKR mice muscles lacking functional IGF1-R, the accumulation of plakoglobin alone can activate insulin signaling (Fig. 5). In any case, plakoglobin seems to enhance insulin potency by triggering intracellular signals, and reduced plakoglobin function is likely to contribute to the development of insulin resistance in type-2 diabetes and DMD^7,8^.

## METHODS

### Animals

All animal experiments were consistent with the ethical guidelines of the Israel Council on Animal Experiments and the Technion Inspection Committee on the Constitution of the Animal Experimentation. Specialized personnel provided animal care in the Institutional Animal facility. We used adult CD-1 male mice (~30g)(Envigo) to isolate plakoglobin containing complexes (Figs. 1 and 2) and study the effects of plakoglobin downregulation (Fig. 3). FVB/N male mice (8-10 weeks old)(Envigo) served as control for the MKR diabetic male mice (8-10 weeks old)(kindly provided by Prof. Derek Le Roithe, Rappaport School of Medicine, Technion Institute, Israel).

### Antibodies and constructs

The plasmids encoding shRNA against plakoglobin and Lacz were previously described^23^. Inserts were cloned into pcDNA 6.2-GW/EmGFP-miR vector using Invitrogen’s BLOCK-iT RNAi expression vector kit. The construct encoding the 6His-tagged plakoglobin has was provided by Dr. Yoshitaka Sekido (Nagoya University School of Medicine, Nagoya, Japan). Plakoglobin antibodies were from Genetex (Cat# GTX15153, lot# 821501500, for immunoprecipitation and immunoblotting) and Acris (Cat# AP33204SU-N, lot# 610051, for immunofluorescence). Anti-Dystrophin (Cat# ab15277, lot# GR 256086-2), anti-Desmin (Cat# ab8592), anti-Vinculin (Cat# ab73412, lot# GR 29133-1), anti-Caveolin-1 (Cat# ab2910, lot# GR 235528-4), anti-Insulin receptor (Cat# ab5500, lot# GR 242593-6, for immunofluorescence), and anti-Myosin (Cat# ab7784) were from Abcam. Anti phospho-insulin receptor (Y1361)(Cat# 3023, lot# 2), anti-Insulin receptor (Cat# 3025, lot# 8, for immunoblotting and immunoprecipitation), anti-phospho-PI3K (Cat# 4228, lot# 2), anti-phospho-AKT (Cat# 9275, lot# 13), anti-Akt (Cat# 9272, lot# 25), and anti-PI3K (Cat# 4292, lot# 9) were from Cell Signaling Technologies. The β-dystroglycan antibody developed by Glenn E. Morris (RJAH Orthopaedic hospital, Oswestry, UK) was obtained from the Developmental Studies Hybridoma Bank, created by the NICHD of the NIH and maintained at The University of Iowa, Department of Biology, Iowa city, IA 52242 (Cat# MANDAG2, clone 7D11, lot# S1ea). Phospho-Serine antibody was from ECM biosciences (Cat# PP2551, lot# 5). Anti-Laminin (Cat# L9393, lot# 046m4837v), and anti-GAPDH (Cat# G8795) were from Sigma. Syntrophin antibody (Cat# Sc-50460, lot# F1109), Caveolin-3 antibody (Cat# sc-5310, lot# L1714), MuSK antibody (clone N-19, Cat# sc-6010), and V5 antibody (Cat# R960-2546-0705, lot# 1831141) were from Santa Cruz Biotechnology (Cat# sc-5310, lot# L1714).

### In vivo transfection

In vivo electroporation was performed as previously described^23^. Briefly, 20μg of plasmid DNA was injected into adult mouse TA muscles, and a mild electric pulse was applied by two electrodes (12V, 5 pulses, 200ms intervals). After 6 days, muscles were dissected and analyzed.

For fiber size analysis, cross-sections of muscles were fixed in 4% PFA and fiber membrane was stained with laminin antibody (see details under immunofluorescence). Cross sectional areas of at least 500 transfected fibers (expressing GFP) and 500 adjacent non-transfected ones in the same muscle section (10µm) were measured using Metamorph (Molecular Devices) and Imaris (Bitplane) software. Images were collected using a Nikon Ni-U upright fluorescence microscope with Plan Fluor 20x 0.5NA objective lens and a Hamamatsu C8484-03 cooled CCD camera and MetaMorph software.

### Isolation of native plakoglobin-containing complexes from skeletal muscle

To isolate plakoglobin-containing complexes from mouse skeletal muscle, we adapted a protocol previously described by Chorev et al., 2014^28^ with some modifications. Briefly, lower limb mouse skeletal muscles (1g) were homogenized in 19 volumes of cold buffer A (20 mM Tris pH 7.2, 100 mM KCl, 5 mM EGTA, 40 mM Imidazole, 1 mM DTT, 1 mM PMSF, 3 mM Benzamidine, 10 μg/ml Leupeptin, 50 mM NaF, 2.7 mM sodium OrthoVanadate, and 1% Triton X-100). Following centrifugation at 10,000 × g for 20 min at 4°C, the pellet (containing myofibrils, membrane and cytoskeleton) was washed once in buffer A, and then resuspended in 10 volumes (v/v) of buffer B (buffer A supplemented with 10 mM ATP) to dissociate myofibrillar myosin from actin thin filaments. After centrifugation at 16,000 × g for 10 min at 4°C, the pellet (mostly containing membrane and cytoskeleton-bound proteins) was washed once in buffer B, resuspended in 10 volumes (v/v) of buffer C (buffer A at pH 9), and incubated at 37°C for 30 min with gentle agitation. After centrifugation at 16,000 × g for 10 min and 4°C, the supernatant (containing purified membrane-bound and cytoskeletal proteins) was subjected to 30% ammonium sulfate precipitation, and precipitates were centrifuged at 20,000 × g for 30 min at 4°C. The obtained pellet was kept on ice and the supernatant was subjected to an additional 50% ammonium sulfate precipitation. After a final centrifugation at 20,000 × g for 30 min at 4°C, precipitates from 30% and 50% ammonium sulfate precipitations were resuspended in 600 ul of buffer A, and 85% of this suspension (containing membrane-cytoskeleton fraction) was loaded onto a Superdex 200 10/300 Gel Filtration column (GE Healthcare Life Sciences, UK) equilibrated with 20 mM Tris pH 7.2. Proteins were eluted at a flow rate of 0.3 ml/min and 300 μl fractions were collected. Selected plakoglobin-containing fractions (Fig. 1d, fractions #8-11) were combined and loaded onto a 1 ml Resource Q anion exchange column (GE Healthcare Life Sciences, UK), equilibrated with 20 mM Tris pH 7.2 and 500 mM NaCl. Proteins were eluted by a 0-500 mM NaCl gradient into 0.5 ml fractions, and fractions enriched with plakoglobin (Fig. 1f lane 7) were analyzed by mass spectrometry, glycerol gradient fractionation and western blotting.

To identify plakoglobin-binding proteins in whole muscle extracts, TA muscles from WT mice (200mg TA from 5 mice) were electroporated with a plasmid encoding 6His-tagged plakoglobin, and 7 d later the muscles were dissected, and whole cell extracts incubated with Ni-beads (according to the manufacturer’s instructions) to purify the 6His-tagged plakoglobin and bound proteins. Interacting proteins were then eluted from Ni-beads using a Histidine gradient (at 50, 80, 100, 150, 200 and 250 mM Histidine), and were analyzed by mass spectrometry, SDS-PAGE and immunoblotting.

### Fractionation of muscle, heart and liver tissues

To obtained whole cell extracts, lower limb muscles (1g) were homogenized as previously described^39^. Briefly, following homogenization in lysis buffer (20 mM Tris pH 7.2, 5 mM EGTA, 100 mM KCl, 1% Triton X-100, 1 mM PMSF, 3 mM Benzamidine, 10 μg/ml Leupeptin, 50 mM NaF, and 2.7 mM Sodium OrthoVanadate), and following centrifugation at 6000 × g for 20 min at 4°C, the supernatant (containing whole cell extract) was collected and stored at −80°C. The pellet was subjected to additional washing steps in homogenization buffer and suspension buffer (20 mM Tris pH 7.2, 100 mM KCl, 1 mM DTT and 1 mM PMSF), and after a final centrifugation at 6000 × g for 20 min at 4°C, the pellet (containing purified myofibrils and desmin filaments) was re-suspended in storage buffer (20 mM Tris pH 7.2, 100 mM KCl, 1 mM DTT and 20% glycerol) and kept at −80°C. To isolate plasma membrane fraction, tissues (muscle, liver, heart) were homogenized in buffer C (20 mM Tris pH 7.6, 100 mM KCl, 5 mM EDTA, 1 mM DTT, 1 mM Sodium OrthoVanadate, 1 mM PMSF, 3 mM Benzamidine, 10 μg/ml Leupeptin, and 50 mM NaF) and centrifuged at 2900 × g for 20 min at 4°C. The obtained supernatant (containing membrane and cytosolic proteins) was then centrifuged at 180,000 × g for 90 min at 4°C using a TLA-55 rotor (Beckman Coulter, Brea, CA). The obtained pellet (containing intact membranes) was then re-suspended in buffer M (20 mM Tris-HCl, pH 7.6, 100 mM KCl, 5mM EDTA, 1mM DTT, 0.25% Sodium deoxycholate, 1% NP-40, 1mM Sodium OrthoVanadate, 10 μg/ml Leupeptin, 3 mM Benzamidine, and 1 mM PMSF), and incubated at 4°C for 20 min with agitation (1400 rpm) to solubilize membrane proteins. After a final centrifugation at 100,000 × g for 30 min at 4°C, the supernatant (i.e. purified membrane fraction) was collected and stored at −80°C.

### Glycerol gradient fractionation

To characterize plakoglobin-containing protein complex, equal amounts of muscle homogenates were layered on a linear 10-40% glycerol gradient containing 20 mM Tris pH 7.6, 5 mM EDTA pH 7.4, 100 mM KCl, 1 mM DTT, 0.24% Sodium Deoxycholate, and 1 mM Sodium Orthovanadate. Following centrifugation at 35,000 rpm for 24 hr at 4°C using a MLS-50 Swinging-Bucket rotor (Beckman Coulter, Brea, CA) to sediment protein complexes, 300μl fractions were collected, and alternate fractions were subjected to Trichloroacetic Acid precipitation (10%) for 24 hr at 4°C. Precipitates were then analyzed by SDS-PAGE and immunoblotting.

### Protein analysis

Immunoblotting and immunoprecipitation were performed as described^23^. In brief, tissue homogenates and fractions of isolated plakoglobin-containing complexes were resolved by SDS-PAGE and immunoblotting with specific antibodies. Immunoprecipitation experiments using specific antibodies (control sample contained mouse IgG) were performed overnight at 4°C, and protein A/G agarose was then added for 4 hr. To remove nonspecific or weakly associated proteins, the resulting precipitates were washed extensively with 10 bed volumes of each of the following buffers: buffer A (50mM Tris-HCl pH 8, 500 mM NaCl, 0.1% SDS, 0.1% Triton, 5 mM EDTA), buffer B (50mM Tris-HCl pH 8, 150 mM NaCl, 0.1% SDS, 0.1% Triton, 5 mM EDTA), and buffer C (50mM Tris-HCl pH 8, 0.1% Triton, 5 mM EDTA). Protein precipitates were eluted in protein loading buffer containing 50 mM DTT and were analyzed by immunoblotting.

### Immunofluorescence labeling of frozen muscle sections

Frozen cross sections of mouse TA were cut at 10 μm or 30 μm (for Z-stack and 3D modeling), fixed in 4% PFA, and incubated in blocking solution (0.2 % BSA and 5 % Normal goat Serum in PBS-T) for 1 hr at RT. Immunofluorescence was performed using plakoglobin (1:50), laminin (1:50), vinculin (1:50), β-dystroglycan (1:30), or insulin receptor (1:30) antibodies overnight at 4°C followed by 1 hr incubation at room temperature with secondary antibodies conjugated to Alexa fluor 568, 647 or DyLight 488 (1:1000). Both primary and secondary antibodies were diluted in blocking solution. Confocal images were collected and analyzed using an inverted LSM 710 laser scanning confocal microscope (Zeiss, Oberkochen, Germany) with a Plan-Apochromat 63 × 1.4 NA objective lens and BP 593/46, BP 525/50 and 640-797 filters, and Imaris 8.2 software. STED images were collected using a Leica DMi8 CS Bino super-resolution microscope (TCS SP8 STED) with a Plan-Apochromat 100 × 1.4 NA oil objective lens, WLL (White Light Laser, covers the spectral range from 470 nm to 670 nm) and the Most Versatile Beamsplitter AOBS, and Leica 3 HyD SP GaAsP-Detector.

PLA assay was performed in humid chamber according to manufacturer’s instructions (Duolink in situ Red, thorugh Sigma). Briefly, muscle cross sections were incubated with β-dystroglycan and insulin receptor antibodies overnight at 4°C, and the control sample was incubated with β-dystroglycan antibody alone. Following incubation with probe-linked secondary antibodies (1 hr at 37°C), the hybridized complementary probes (those that are within 20-100nm) were ligated (30 min at 37°C) to form circular DNA. Finally, rolling circle amplification reaction and incorporation of red fluorescently labeled nucleotides were allowed to proceed for 3 hr at 37°C. Red fluorescent spots were visualized and images collected using an inverted LSM 710 laser scanning confocal microscope (Zeiss, Oberkochen, Germany) with a Plan-Apochromat 63 × 1.4 NA objective lens and BP 593/46 filter, and Imaris 8.2 software.

### [^3^H]-2-deoxyglucose uptake by skeletal muscle

For *ex-vivo* glucose uptake by skeletal muscle, we used a modified protocol adapted from Fernandez et al ^4^, Lauro et al ^56^, and Witczak et al ^57^. Mice were fasted for 12 hr and euthanized with CO_2_. TA muscles were carefully removed, weighted, and incubated for 2 hr in Krebs-Ringer bicarbonate (KRB) buffer (117 mM NaCl, 4.7 mM KCl, 2.5 mM CaCl_2_, 1.2 mM MgSO_4_, 1.2 mM KH_2_PO_4_ and 25 mM NaHCO_3_), which was pre-equiliberated to 95% O_2_ and 5% CO_2_ and adjusted to 37°C. To measure glucose uptake, dissected muscles were transferred to KRB buffer containing 1.5 μCi/ml of [^3^H]-2-deoxyglucose, 200 nM insulin (Sigma), 0.1 mM 2-deoxy-D-glucose (Sigma) and 7mM Manitol (J.T.Baker) for 45 min at 95% O_2_, 5% CO_2_ and 37°C. Cytochalasin B (20 nM), a glucose transport inhibitor (Sigma), was added to control tubes. Then, muscles were quickly blotted on a filter paper and frozen in liquid nitrogen to stop the reaction. To measure glucose uptake, muscles were homogenized in 500 μl lysis buffer (20 mM Tris pH 7.4, 5 mM EDTA, 10 mM Sodium pyrophosphate, 100 mM NaF, 2 mM Sodium OrthoVanadate, 10 μg/ml Aprotinin, 10 μg/ml Leupeptin, 3 mM Benzamidine and 1 mM PMSF) and rates of [^3^H]-2-deoxyglucose uptake were assessed in muscle homogenates using scintillation counter as described^57^.

For glucose bolus experiments, D-glucose (1 mg/gr body weight) was administered to mice by retro-orbital injection. TA muscles were collected 25 min after injection and kept frozen for further analysis.

### Mass Spectrometry analysis

To identify the components comprising the plakoglobin-containing multiprotein assemblies, fractions eluted as a high molecular mass peak from the gel filtration column (Fig. 1c), as a sharp peak from the anion exchange column (Fig. 1f lane 7), or as 6His-tagged plakoglobin bound proteins (Fig. 2a) were analyzed by SDS-PAGE and coomassie blue staining. The stained protein bands were reduced with 3 mM DTT at 60ºC for 30 min, modified with 10 mM iodoacetamide in 100 mM ammonium bicarbonate in the dark at room temperature for 30 min, and then digested in 10% acetonitrile and 10 mM ammonium bicabonate with modified trypsin (Promega) at a 1:10 enzyme-to-substrate ratio overnight at 37°C. The tryptic peptides were desalted using C18 tips (Homemade stage tips), dried, and re-suspended in 0.1% Formic acid. The peptides were resolved by reverse-phase chromatography on 0.075 × 180-mm fused silica capillaries (J&W) and packed with Reprosil reversed phase material (Dr Maisch GmbH, Germany). The peptides were eluted with linear 60 min gradient of 5 to 28%, 15 min gradient of 28 to 95%, and 15 min at 95% acetonitrile with 0.1% formic acid in water at flow rates of 0.15 μl/min. Mass spectrometry was performed by Q Exactive plus mass spectrometer (Thermo) in a positive mode using repetitively full MS scan followed by collision induces dissociation (HCD) of the 10 most dominant ions selected from the first MS scan. The mass spectrometry data was analyzed using Proteome Discoverer 1.4 software with Sequest (Thermo), and Mascot (Matrix Science) algorithms against mouse Uniprot database. Results were filtered with 1% false discovery rate. Semi quantitation was done by calculating the peak area of each peptide based on its extracted ion currents (XICs), and the area of the protein is the average of the three most intense peptides from each protein.

### Statistical analysis and image acquisition

Data are presented as means and error bars indicate SEM. The statistical significance was accessed by one-tailed Student’s t-test. Images were processed by Adobe Photoshop CS5, version 12.1. Quantity One algorithm (Bio-Rad Laboratories version 29.0) was used for densitometric measurements of protein bands intensity.

Confocal images were collected and analyzed using an inverted LSM 710 laser scanning confocal microscope (Zeiss, Oberkochen, Germany) with a Plan-Apochromat 63 × 1.4 NA objective lens and BP 593/46, BP 525/50 and 640-797 filters, and Imaris 8.2 software. STED images were collected using a Leica DMi8 CS Bino super-resolution microscope (TCS SP8 STED) with a Plan-Apochromat 100 × 1.4 NA oil objective lens, WLL (White Light Laser, covers the spectral range from 470 nm to 670 nm) and the Most Versatile Beamsplitter AOBS, and Leica 3 HyD SP GaAsP-Detector. STED images were analyzed using the STED image reconstruction module of Leica. Sub-diffraction spots colocalization analysis and visualizations was implemented using Imaris 9.1.2 software (Oxford, Bitplane). IMARIS spots module was set to identify sub-diffraction spots as particles with 150nm diameter and quality threshold value above 3000; to identify triple spots co-occurrence, the insulin receptor staining was used as baseline to filter-in the other proteins beyond intensity threshold value of 1500, and to refine the positioning of the triple particles. Moreover, a built-in Imaris plugin spot colocalization algorithm (requiring ImarisXT module) was used to identify triple spots centers with maximum distance of 250nm.

## Data availability

The datasets generated and analysed during the current study are available from the corresponding author on request.

## ACKNOWLEDGMENTS

This project was supported by grants from the Israel Science Foundation (grant No. 623/15) to S. Cohen. Additional funds have been received from The Russell Berrie Nanotechnology Institute, Technion.

We thank Prof. Derek Leroith from Rambam medical center for kindly sharing the MKR mice with us. We also thank Dr. Dror Chorev and Prof. Michal Sharon from Weizmann institute for their assistance with the protocol for the tandem native purification approach. We are grateful for the kind assistance we received from Dr. Carol Witczak with the glucose uptake approach. We acknowledge the Smoler Proteomics Center at the Technion for their assistance with the proteomics work, the LS&E infrastructure unit for their help with the confocal microscopy, and the microscopy center at Bar Ilan University (Ramat-Gan and Safed) for their assistance with the super-resolution STED microscopy.

