## Supplementary Tables for "Novel signaling hub of insulin receptor, dystrophin glycoprotein complex and plakoglobin regulates muscle size"

**Table S1. Membrane, cytoskeletal, and myofibrillar components that were co-purified with plakoglobin from skeletal muscle.**

| **Protein name** | **Gene** | **# Unique peptides** |
|---|---|---|
| Plakoglobin | *JUP* | 15 |
| Sarcoglycan, delta | *SGCD* | 3 |
| Laminin, beta 2 | *LAMB2* | 4 |
| Laminin, gamma 1 | *LAMC1* | 17 |
| Laminin Subunit Beta 1 | *LAMB1* | 21 |
| Spectrin alpha 2 | *SPNA2* | 7 |
| Spectrin beta 1 | *SPNB1* | 8 |
| Desmin | *DES* | 9 |
| Plectin | *PLEC* | 36 |
| Actinin, alpha 1 | *ACTN1* | 23 |
| Actinin alpha 2 | *ACTN2* | 29 |
| Actinin alpha 3 | *ACTN3* | 77 |
| Actinin alpha 4 | *ACTN4* | 12 |
| Plakophilin 1 | *PKP1* | 2 |
| Desmoglein 1 | *DSG1C* | 2 |
| Desmoplakin | *DSP* | 25 |
| Cadherin 13 | *CDH13* | 2 |

Native protein assemblies of plakoglobin and bound proteins were isolated from membrane-cytoskeletal preparations of lower limb mouse muscles by size exclusion chromatography and anion-exchange column. Components that were co-purified with plakoglobin as a major protein peak were identified by mass spectrometry.

**Table S2. Plakoglobin binds membrane, cytoskeletal, myofibrillar and insulin signaling components in skeletal muscle homogenates (6,000g supernatant).**

| Protein name | Gene (NCBI) | # Unique peptides | Present in high MW peak |
|---|---|---|---|
| Plakoglobin | *JUP* | 16 | + |
| Dystrophin | *DMD* | 20 | |
| Sarcoglycan, alpha | *SGCA* | 3 | |
| Sarcoglycan, beta | *SGCB* | 2 | |
| Sarcoglycan, delta | *SGCD* | 2 | + |
| Alpha-1-syntrophin | *SNTA1* | 9 | |
| Beta-2-syntrophin | *SNTB2* | 4 | |
| Dystroglycan | *DAG1* | 3 | |
| Nitric oxide synthase 1 | *NOS1* | 3 | |
| Vinculin | *VCL* | 6 | |
| Caveolin 1 | *CAV1* | 5 | |
| Caveolin 3 | *CAV3* | 2 | |
| Laminin, beta 2 | *LAMB2* | 3 | + |
| Laminin, gamma 1 | *LAMC1* | 2 | + |
| Spectrin alpha 2 | *SPNA2* | 26 | + |
| Spectrin beta 1 | *SPNB1* | 13 | + |
| Insulin receptor substrate 1 | *IRS1* | 7 | |
| Insulin-like growth factor 2 receptor | *IGF2R* | 9 | |
| Insulin-like growth factor binding protein | *IGFALS* | 6 | |
| PI3K Catalytic Subunit Type 3 | *PIK3C3* | 1 | |
| PI3K Catalytic Subunit Gamma | *PIK3CG* | 2 | |
| PI3K-p85 | *PIK3R1* | 4 | |
| Desmin | *DES* | 26 | + |
| Plectin | *PLEC* | 147 | + |
| Actinin, alpha 1 | *ACTN1* | 6 | + |
| Actinin alpha 2 | *ACTN2* | 1 | + |
| Actinin alpha 3 | *ACTN3* | 2 | + |
| Actinin alpha 4 | *ACTN4* | 12 | + |
| Cadherin 13 | *CDH13* | 6 | + |

Affinity purification of 6His-tagged plakoglobin and associated proteins from the soluble fraction of mouse lower limb muscles. Mass spectrometry analysis identified membrane, cytoskeletal and myofibrillar components (also present in the high MW peak, see Table I), as well as component of PI3K-Akt signaling.
