## Supplementary figures and images for "Novel signaling hub of insulin receptor, dystrophin glycoprotein complex and plakoglobin regulates muscle size"

### Figure S1

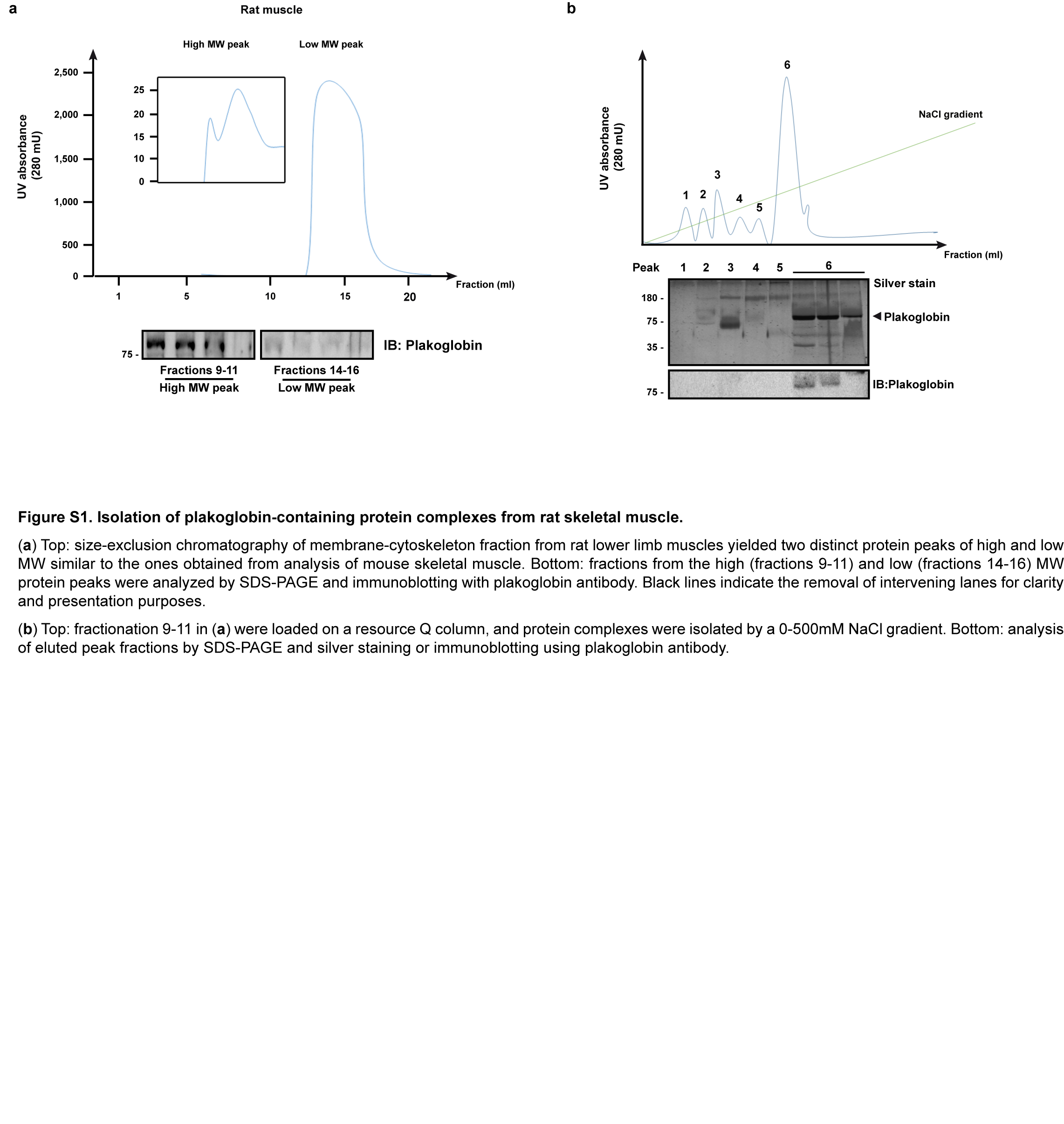

### Figure S2

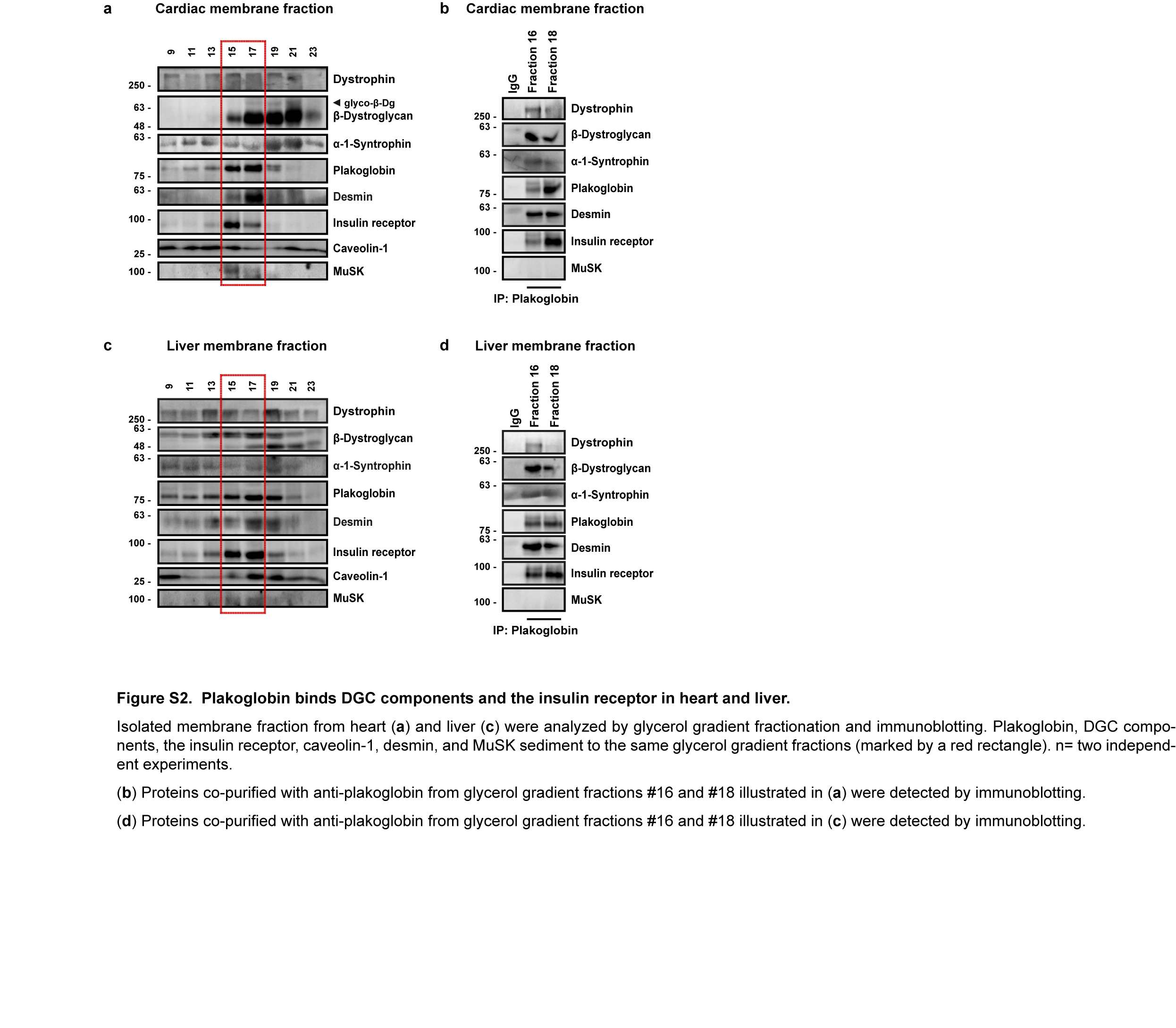

### Figure S3

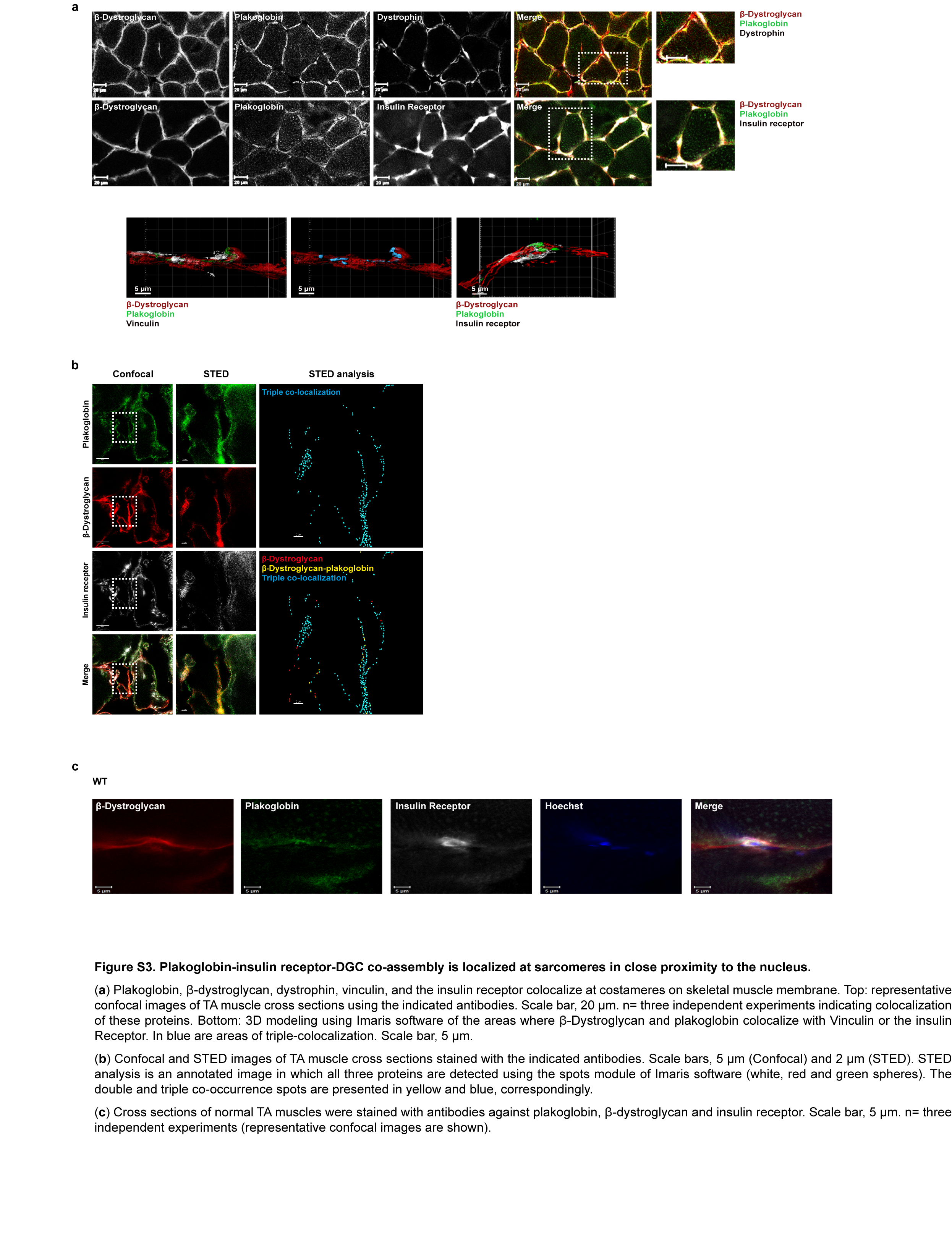

### Figure S4

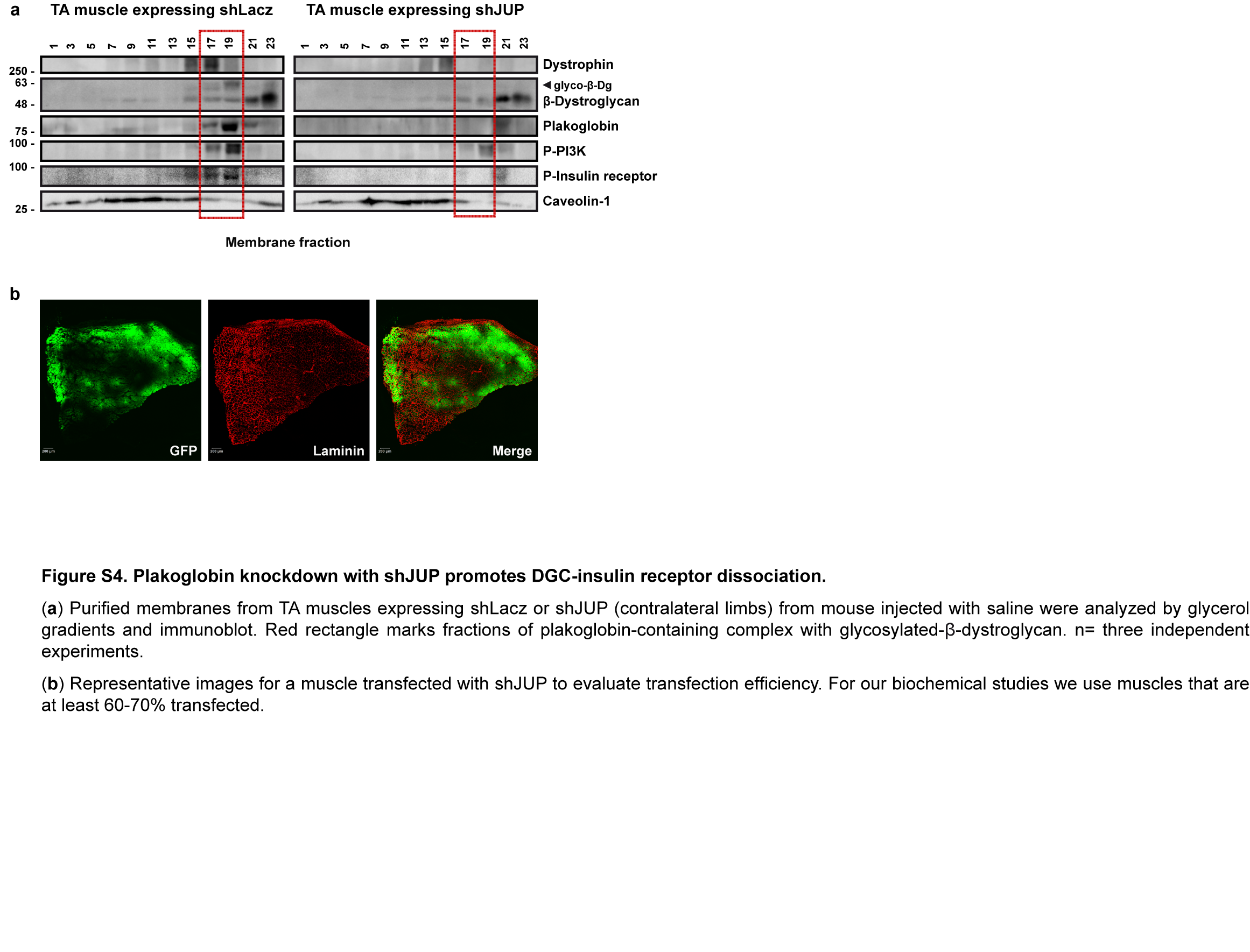

### Figure S5

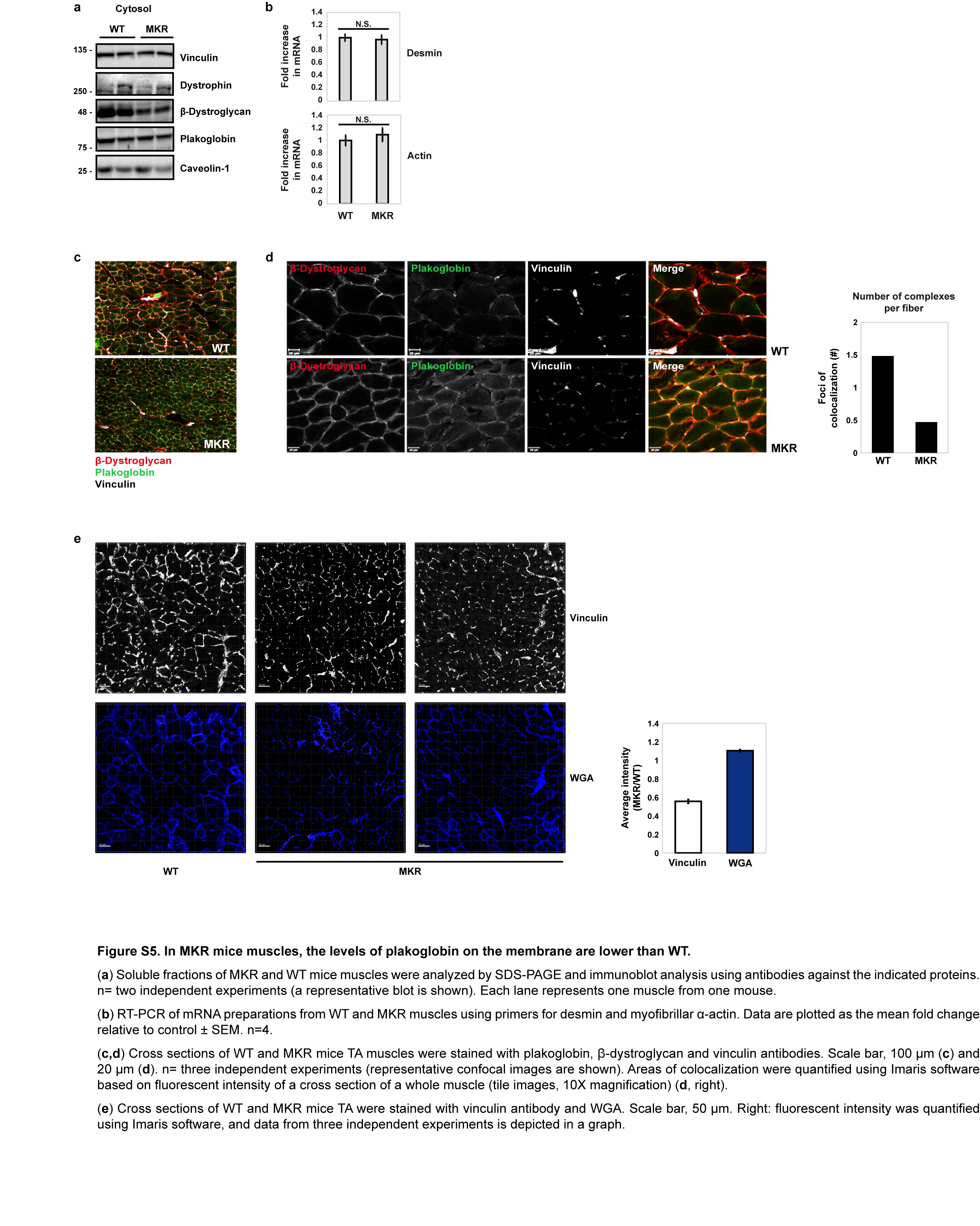
